# Rho1 activation recapitulates early gastrulation events in the ventral, but not dorsal, epithelium of *Drosophila* embryos

**DOI:** 10.1101/2020.03.12.989285

**Authors:** Ashley Rich, Richard G. Fehon, Michael Glotzer

## Abstract

Ventral furrow formation, the first step in *Drosophila* gastrulation, is a well-studied example of tissue morphogenesis. Rho1 is highly active in a subset of ventral cells and is required for this morphogenetic event. However, it is unclear whether spatially patterned Rho1 activity alone is sufficient to recapitulate all aspects of this morphogenetic event, including anisotropic apical constriction and coordinated cell movements. Here, using an optogenetic probe that rapidly and robustly activates Rho1 in *Drosophila* tissues, we show that Rho1 activity induces ectopic deformations in the dorsal and ventral epithelia of *Drosophila* embryos. These perturbations reveal substantial differences in how ventral and dorsal cells, both within and outside the zone of Rho1 activation, respond to spatially and temporally identical patterns of Rho1 activation. Our results demonstrate that an asymmetric zone of Rho1 activity is not sufficient to recapitulate ventral furrow formation and indicate that additional, ventral-specific factors contribute to the cell- and tissue-level behaviors that emerge during ventral furrow formation.

## Introduction

Tissue morphogenesis underlies the development of multicellular organisms. The molecular and cellular mechanisms that govern tissue morphogenesis remain a central challenge in developmental cell biology. Extensive genetic and biochemical experiments have defined the factors required for many morphogenetic movements. Furthermore, methods for imaging and quantitatively describing cell shape changes are ever-improving. Despite this progress, questions remain. For example, how pliable are tissues before and while they are deforming? To what degree does the underlying cytoskeleton of cells within a tissue limit their ability to deform, and to what degree are shape changes of neighboring cells coordinated?

Ventral furrow formation in the *Drosophila* embryo is one of the best studied examples of tissue morphogenesis; it is the first step in *Drosophila* gastrulation. Ventral furrow formation occurs when a rectangular zone of approximately 1000 cells, arranged in 18 rows, on the ventral surface of the embryonic epithelium apically constrict and invaginate into the embryo, ultimately giving rise to the embryonic mesoderm (***Leptin and Grunewald, 1990***; ***Sweeton et al., 1991***). Many molecules required for ventral furrow formation have been identified: An extracellular serine protease cascade activates the transcription factor Dorsal which drives the expression of two additional transcription factors, Snail and Twist, in a subset of ventral cells, inducing them to adopt mesodermal fates (***Morisato and Anderson, 1995***; ***Ip et al., 1992***; ***Jiang et al., 1991***). Snail and Twist then induce the expression of secreted and cell surface molecules, including the ligand Fog, the G-protein-coupled receptor (GPCR) Mist, and the transmembrane protein T48 (***Dawes-Hoang et al., 2005***; ***Costa et al., 1994***; ***Kölsch et al., 2007***; ***Manning et al., 2013***). Together with Concertina, a maternally contributed G*α* protein, and Smog, a maternally contributed GPCR, these factors recruit and activate RhoGEF2, a Rho1-specific guanine nucleotide exchange factor, at the apical membrane of ventral cells (***Parks and Wieschaus, 1991***; ***Kölsch et al., 2007***; ***Nikolaidou and Barrett, 2004***; ***Kerridge et al., 2016***). RhoGEF2 then activates Rho1 to assemble a contractile actomyosin network (***Fox and Peifer, 2007***); these networks within single cells are coupled through adherens junctions between neighboring cells into a supracellular actomyosin network that promotes apical constriction and robust ventral furrow formation (***Martin et al., 2010***; ***Yevick et al., 2019***). Notably, both RhoGEF2 accumulation and Rho1 activation are pulsatile (***Martin et al., 2010***; ***Mason et al., 2016***).

The intracellular signaling cascade described above activates Rho1 within individual presumptive mesoderm cells. This could, in principle, account for ventral furrow formation (***Gilmour et al., 2017***; ***Ko and Martin, 2020***). However, several features of the ventral furrow suggest that ventral cells exhibit a high degree of intercellular coupling, which may influence the outcome of the genetically encoded contractility. For example, the cells in the ventral furrow constrict their apices more along the dorsal-ventral axis of the embryo than along the anterior-posterior axis (***Sweeton et al., 1991***; ***Martin et al., 2010***). If individual ventral cells constrict and invaginate without being influenced by their neighbors, one would predict isotropic apical constriction. Additionally, the apical constriction of individual cells appears coordinated, with cells adjacent to constricting cells more likely to constrict than their more distant counterparts (***Sweeton et al., 1991***; ***Gao et al., 2016***). Furthermore, multiple rows of cells lateral to the furrow bend towards it, indicating that forces are transmitted over long distances in the ventral epithelium (***Rauzi et al., 2015***; ***Costa et al., 1994***; ***Leptin et al., 1992***).

Taken together, this wealth of previous results suggests that ventral furrow formation results from a combination of intracellular Rho1-mediated contractility and intercellular coupling of those contractile forces. In the simplest iteration of this model, an asymmetric zone of Rho1 activation is sufficient to recapitulate both the intra- and intercellular aspects of ventral furrow formation (***Doubrovinski et al., 2018***). Indeed, it was recently shown that an asymmetric zone of local Rho1 activation is sufficient to induce an ectopic invagination in the dorsal *Drosophila* epithelium (***Izquierdo et al., 2018***). However, it remains unclear whether local Rho1 activation alone is sufficient to induce sustained tissue morphogenesis and recapitulate all aspects of ventral furrow formation, or whether furrows in wildtype embryos result from a local zone of contractility modulated by ventral-specific gene expression.

Addressing these and related questions necessitates the ability to activate Rho1 with high spatial and temporal precision without otherwise perturbing the embryo. Optogenetic techniques utilize photosensitive proteins to control protein localization and/or activity with light; these techniques are, therefore, well-suited to interrogate the basis for the anisotropic and coordinated nature of apical constriction during ventral furrow formation. Importantly, the ideal optogenetic approach will activate Rho1 in response to light alone.

Here, we use a LOV-domain based optogenetic probe to acutely activate Rho1 in *Drosophila*. We demonstrate that this system expresses ubiquitously throughout *Drosophila* development and is well tolerated. Optogenetic activation of Rho1 induces ectopic deformations in both the dorsal and ventral embryonic epithelium at the onset of gastrulation. We find that ventral cells specifically respond to ectopic Rho1 activation with aligned, anisotropic apical constriction. This ventral-specific response requires Dorsal and Twist expression. Furthermore, we provide evidence that the transmission of contractile forces over long distances is specific to the ventral epithelium.

## Results

### A LOV domain-based optogenetic system controls Rho1 activity in *Drosophila*

To study the cellular consequences of acute Rho1 activation and probe the impact of Rho1 activation on cells neighboring the activation region, we adapted an optogenetic system for use in *Drosophila*. This two-component system consists of a membrane tethered LOV domain fused to the SsrA peptide and a cytoplasmic SspB protein fused to a protein of interest (***Figure 1a***) (***Guntas et al., 2015***; ***Strickland et al., 2012***). Blue light exposure induces a conformational change in the LOV domain, exposing the SsrA peptide and recruiting the SspB fusion protein to the plasma membrane (***Figure 1a***). As a proof of concept, we first expressed the membrane localized LOV domain and an SspB-mScarlet fusion from the ubiquitin promoter. Local activation of a region of the dorsal embryonic epithelium with blue light induces rapid recruitment of SspB-mScarlet to the plasma membrane (***Figure 1b***). SspB-mScarlet remains associated with the plasma membrane as long as blue light activation is sustained but rapidly (∼ 1 min) returns to its dark state, cytoplasmic localization, upon cessation of photoactivation (***Figure 1b***).

**Figure 1.**
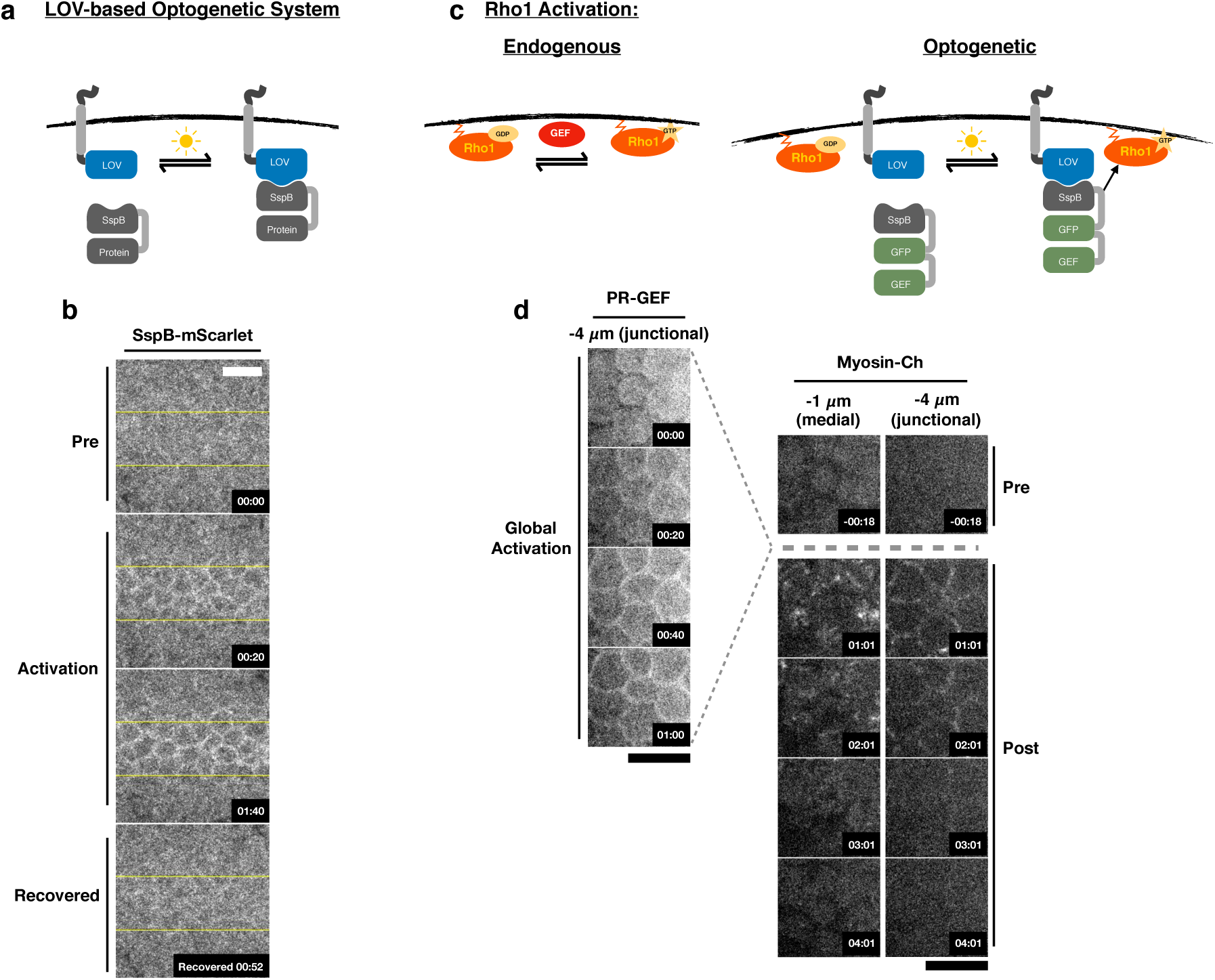
Optogenetic control of Rho1 in *Drosophila*. a) Generic LOV domain-based optogenetic system consisting of a membrane localized LOVSsrA protein and a recruitable SspB protein. SspB can be fused to any protein of interest. Blue light induces a conformational change in the LOV domain, allowing it to recruit SspB fusion proteins to the membrane. b) Dorsal epithelium of an embryo expressing the membrane-localized LOVSsrA and SspB-mScarlet at the onset of gastrulation before, during, and after photo-activation in the indicated region (yellow box). Data representative of 4/4 embryos. c) Optogenetic system for activating Rho1: SspB is fused to GFP and the Dbl homology (DH) domain of LARG (PR-GEF). Photoactivation induces recruitment of the PR-GEF to the membrane and Rho1 activation. d) Dorsal epithelium of an embryo expressing a membrane-localized LOVSsrA and PR-GEF at the onset of gastrulation. The distribution of PR-GEF and myosin are shown before, during, and at the indicated times after global photoactivation. Data representative of 5/5 embryos. Time zero indicates the beginning of photoactivation. Scale bars are 10 μm. **Figure 1–Figure supplement 1.** Recruitment of PR-GEF activates Rho1 in all tissues tested. **Figure 1–Figure supplement 2.** Ectopic Rho1 activation is sensitive to light dose. **Figure 1–Figure supplement 3.** Inactivation kinetics of the LOV domain dictate the off rate of optogenetic-induced Rho1 activity.

To control Rho1 activation, we replaced mScarlet with the catalytic Dbl homology (DH) domain of LARG to generate photorecruitable SspB-GFP-LARG(DH) (hereafter called PR-GEF) (***Figure 1c***). LARG is a human RhoA-specific GEF; the DH domain of LARG has previously been used in a related optogenetic system to control RhoA activity in mammalian tissue culture cells (***Wagner and Glotzer, 2016***). We used the DH domain of LARG alone to ensure that the recruitable GEF’s function is divorced from all endogenous regulation and only sensitive to optogenetic activation. Homozygous flies expressing the membrane localized LOVSsrA and PR-GEF from the ubiquitin promoter are viable and fertile, indicating that these transgenes are well tolerated. Global activation of the dorsal embryonic epithelium with blue light induces strong recruitment of PR-GEF to the plasma membrane within seconds, and this global PR-GEF recruitment induces cortical myosin accumulation within 1 minute (***Figure 1d***). Myosin accumulates both medially and junctionally (***Figure 1d***). Optogenetically-induced cortical myosin completely disappears within 3 minutes of cessation of photoactivation (***Figure 1d***). Thus, using conventional instruments, this optogenetic system rapidly, robustly, and reversibly activates Rho1 in the embryonic epithelium. This system also activates Rho1 in all *Drosophila* tissues tested, including the pupal notum, follicular epithelium, larval wing imaginal disc, and larval central nervous system (***Figure 1–Figure Supplement 1***). In the larval wing peripodial epithelium, optogenetic activation of Rho1 can induce myosin accumulation with subcellular precision (***Figure 1–Figure Supplement 1c***). Optogenetic activation of Rho1 is sensitive to light dosage; attenuating the activating light induced less myosin-Ch accumulation, indicating lower levels of Rho1 activation (***Figure 1–Figure Supplement 2***). Above a certain threshold of activating light, Rho1 becomes globally activated, despite precisely defined activation regions (***Figure 1–Figure Supplement 2***). Thus, this LOV domain-based optogenetic probe is capable of controlling Rho1 activation with high spatial and temporal resolution. Furthermore, the level of Rho1 activation can be tuned by modulating light dosage.

While this LOV domain-based optogenetic probe recovers to its dark state activity level within minutes (***Figure 1***), some biological phenomena may require faster recovery kinetics. To increase the inactivation rate of our optogenetic probe, we introduced a previously identified point mutation, I427V, into the LOV domain, which increases the rate at which the LOV domain returns to the dark state (***Christie et al., 2007***). I427V increases the inactivation rate of the optogenetic system in *Drosophila* S2 cells expressing a membrane localized LOV domain containing this mutation and a cytoplasmic, recruitable tagRFP-SspB (***Figure 1–Figure Supplement 3a,b***). Increasing the recovery rate of the LOV domain also decreases the maximum recruitment of tagRFP-SspB (***Figure 1–Figure Supplement 3a,b***), demonstrating the trade off between rapid inactivation and total recruitment. Global activation of the rapid cycling LOV domain in *Drosophila* larval brains induced robust Rho1 activation, as scored by accumulation of a Rho1 biosensor (***Figure 1–Figure Supplement 3c***); this Rho1 activity dissipated within a minute of cessation of global optogenetic activation. In contrast, Rho1 remained active a minute after global activation of the wild type LOV domain (***Figure 1–Figure Supplement 3c***). Thus, the cycling kinetics of the LOV domain are the primary determinant of the off rate of optogenetic-induced Rho1 activity. This emphasizes that there are rapid and robust mechanisms for shutting off Rho1 activity in vivo; furthermore, it suggests that cells continually activate Rho1 during cellular and developmental processes that require sustained Rho1 activation. The wildtype LOV domain is used for the remainder of the experiments presented, as the rapid recovery was not essential to address the questions answered here.

### Rho1 activation is sufficient to induce reversible invaginations in the *Drosophila* embryonic epithelium

After validating that this system reversibly activates Rho1 in the early *Drosophila* embryo, we asked whether an asymmetric zone of Rho1 activation is sufficient to induce an ectopic invagination in the embryonic dorsal epithelium just after cellularization. Ectopic Rho1 activation induces apical myosin accumulation and is sufficient to induce an ectopic invagination (***Figure 2a***). Importantly, the size of the invaginated region closely mirrors the size of the photoactivated zone, demonstrating the spatial precision of this approach and emphasizing that this deformation is light-dependent. (Rho1 is activated in asymmetric zones of the same dimensions throughout this work, except where explicitly stated.) This is consistent with recently published work (***Izquierdo et al., 2018***).

**Figure 2.**
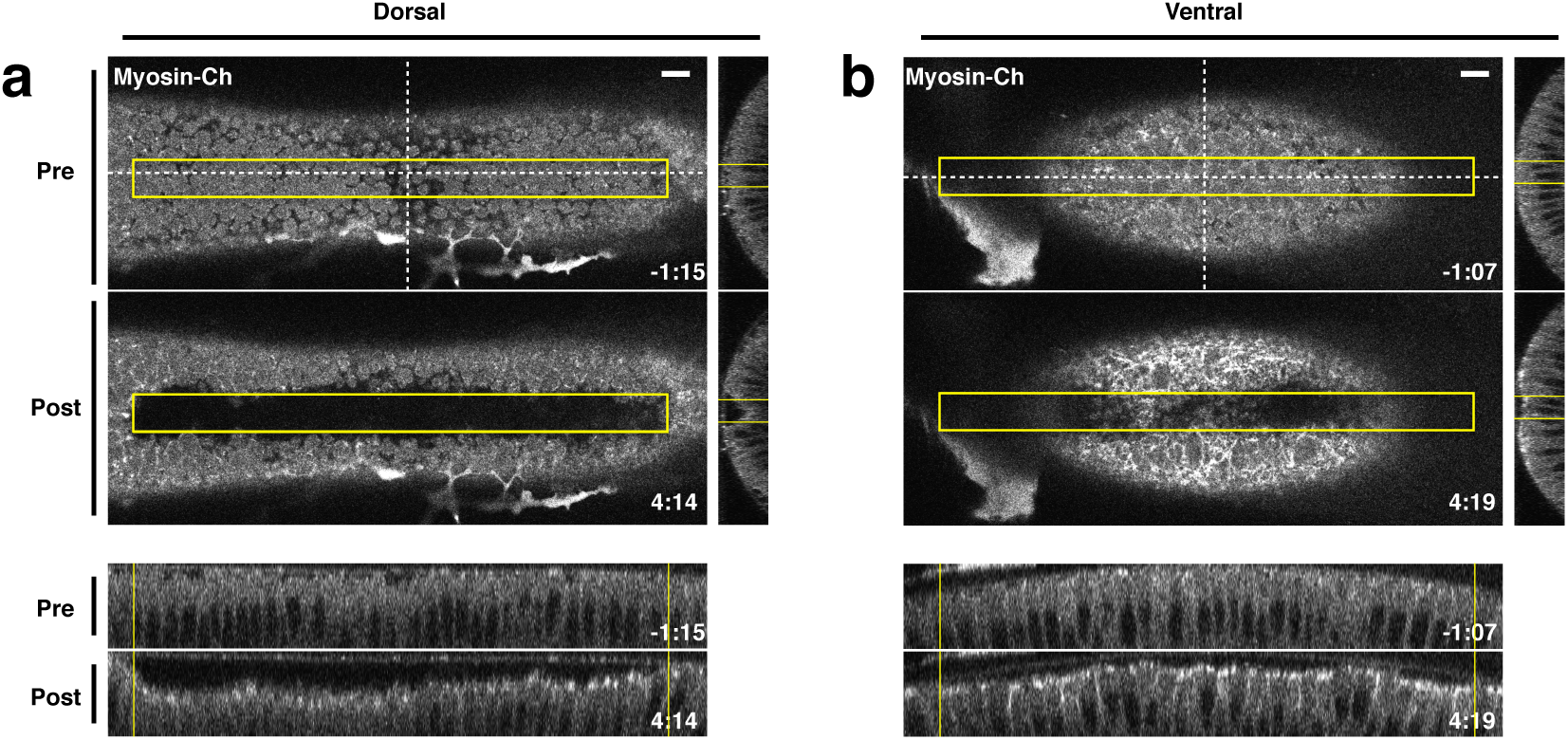
Local Rho1 activation is sufficient to induce ectopic invaginations in the embryonic epithelium. a,b) Embryos expressing the optogenetic components and Myosin-Ch at the onset of gastrulation were optogenetically activated in a single apical plane within the yellow box. Ectopic invaginations and Myosin-Ch accumulation are shown in the dorsal (a) and ventral (b) epithelium. Myosin-Ch accumulates both within and outside the region of optogenetic Rho1 activation in the ventral epithelium (b), YZ projection, Right; XZ project, Bottom). This is due to endogenous gastrulation. Data representative of 7/7 (a) and 5/5 (b) embryos. Time zero indicates the first pulse of blue light activation. Scale bars are 10 μm. **Figure 2–Figure supplement 1.** Optogenetic-induced invaginations revert following cessation of Rho1 activation.

Local Rho1 activation also induces ectopic invaginations in the ventral embryonic epithelium prior to the onset of ventral furrowing (***Figure 2b***). The ectopic invaginations induced in either the dorsal or ventral epithelium recover to their pre-activation state within four minutes of cessation of optogenetic Rho1 activation (***Figure 2–Figure Supplement 1***). These recovery kinetics are slightly longer than the cycling kinetics of the WT LOV domain (***Figure 1d***, ***Figure 1–Figure Supplement 3***). Thus, while ectopic Rho1 activation induces ectopic deformations, the downstream consequences of optogenetic Rho1 activation are rapidly and robustly inactivated in the absence of continued photoactivation.

### Rho1 activation induces distinct apical constriction in the dorsal and ventral epithelium

To determine whether dorsal and ventral cells respond similarly to spatially and temporally identical zones of Rho1 activation, we locally activated Rho1 in the dorsal or ventral epithelium of embryos expressing the optogenetic components and the membrane marker Gap43-mCh and subsequently segmented tissues, tracked individual cells, and quantified the area of the apical-most surface of dorsal or ventral cells before and after optogenetic Rho1 activation (***Aigouy et al., 2010***). We also quantified the anisotropy of apical constriction by measuring the extent of elongation of the apical-most surfaces of cells before and after photoactivation (***Aigouy et al., 2010***). Local Rho1 activation in individual cells in the dorsal embryonic epithelium induced apical constriction (***Figure 3a***); optogenetic activation of Rho1 in a collection of dorsal cells also induced apical constriction (***Figure 3b-c***). Unlike cells of the endogenous ventral furrow, optogenetically activated dorsal cells constrict isotropically (***Figure 3d***-bottom panel, ***Figure 3–Figure Supplement 4***). Thus, an asymmetric, rectangular zone of Rho1 activation via a LOV-domain based probe is not sufficient to fully recapitulate the cell shape changes associated with endogenous ventral furrowing. This result differs from previous work (See Discussion) (***Izquierdo et al., 2018***).

**Figure 3.**
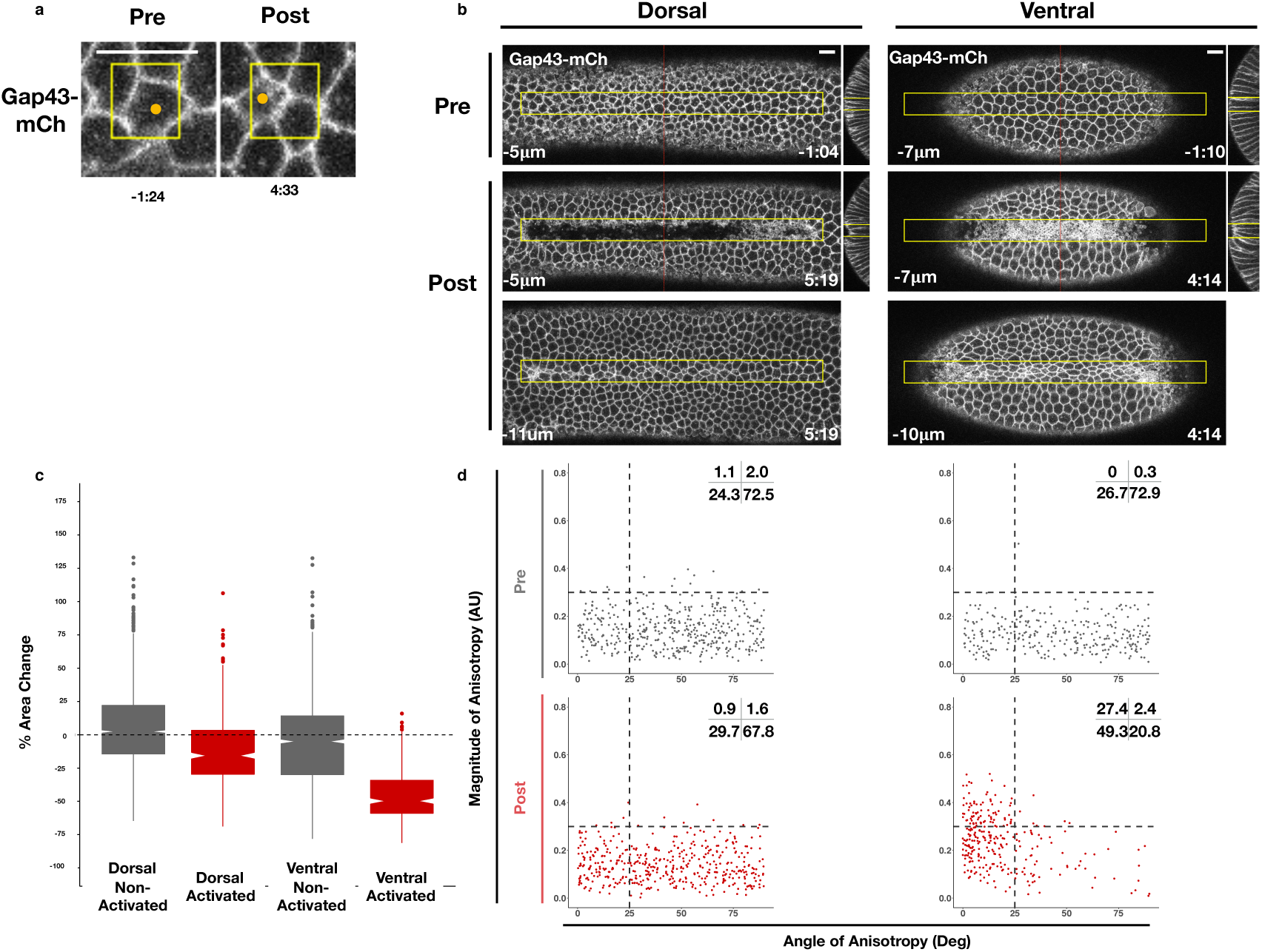
Rho1 activation induces distinct apical constriction in dorsal and ventral epithelial cells. a) Dorsal cells from an embryo expressing the optogenetic components and Gap43-mCh at the onset of gastrulation before and after photoactivation within the yellow box. Z-slices shown represent the apical-most view of the activated cell (orange dot). Data representative of 7/8 cells from 2 embryos. b) Dorsal (left) and ventral (right) epithelium of embryos expressing the optogenetic components and Gap43-mCh at the onset of gastrulation. Rho1 was activated in the yellow box. Images shown in bottom panels were chosen to show the apical surfaces of activated cells. Red lines in (b) indicate position of each YZ slice. Data are representative of 4/4 (a) and 4/4 (b) embryos. Time zero indicates the first pulse of blue light activation. Scale bars are 10 μm. c) Quantification of apical area change induced by optogenetic Rho1 activation. Gray columns represent non-activated cells (cells outside the yellow box in (b)). Red columns represent activated cells (cells within the yellow box in (b)). d) Anisotropy scatter plots: Each dot represents an activated dorsal (left) or ventral (right) cell before (top) or after (bottom) optogenetic activation of Rho1. The magnitude of anisotropy is plotted on the y-axis; the orientation of anisotropy, relative to the anterior-posterior axis of the embryo, is plotted on the x-axis. Dotted lines are provided to facilitate comparisons. Insets show percentage of cells in each quadrant. Cells in the upper left quadrant exhibit highly aligned, anisotropic apical constriction. 444 dorsal cells from 4 embryos and 288 ventral cells from 4 embryos were analyzed. See ***Figure 3–Figure Supplement 4*** for plots of the changes in anisotropy of individual cells. **Figure 3–Figure supplement 1.** Quantification of endogenous ventral furrow formation. **Figure 3–Figure supplement 2.** Schematic of data collection and analysis for local activation experiments. **Figure 3–Figure supplement 3.** Optogenetic activation of Rho1 induces precocious cell shape changes in the ventral epithelium. **Figure 3–Figure supplement 4.** WT ventral cells exhibit large changes in the magnitude and alignment of anisotropy in response to Rho1 activation.

We next tested whether optogenetic Rho1 activation has a different effect on ventral cells, which express ventral-specific genes. We activated ventral cells before they exhibited any overt signs of apical constriction. Optogenetic activation of Rho1 in a collection of ventral cells also induced apical constriction within the zone of Rho1 activation (***Figure 3b,c***). However, the apical surfaces of activated ventral cells were elongated, indicating anisotropic apical constriction. This anisotropy was strongly aligned with the anterior-posterior axis (***Figure 3d***, ***Figure 3–Figure Supplement 4***). A smaller, asymmetric zone of Rho1 activation also induces anisotropic apical constriction in activated ventral cells but not their non-activated neighbors (***Figure 3–Figure Supplement 3***), confirming that our optogenetic experiments induce precocious cell shape changes in the ventral epithelium. In contrast to the isotropic apical constriction induced in activated cells within the dorsal epithelium, optogenetic Rho1 activation in cells within the ventral epithelium induces precocious, anisotropic apical constriction that strongly resemble the anisotropic apical constrictions seen during endogenous ventral furrow formation (***Figure 3–Figure Supplement 1***).

### Genetic requirements for ventral-specific responses

The finding that an asymmetric zone of Rho1 activation induces differential responses in the dorsal and ventral epithelia suggests that ventral patterning may influence the response to Rho1 activation. Ventral-specific factors, such as those downstream of Dorsal, Twist, and/or Snail, may cooperate with an asymmetric zone of Rho1 activation during endogenous ventral furrow formation to drive strong, anisotropic apical constriction. To test this hypothesis, we locally activated Rho1 in ventral cells lacking Dorsal protein, a factor required for ventral identity. Optogenetic Rho1 activation in embryos derived from females homozygous for a null *dorsal* allele still induced apical constriction, but these apical constrictions were weaker than those of WT ventral cells and were no longer anisotropic (***Figure 4, Figure 3–Figure Supplement 4***). Indeed, in the absence of the Dorsal protein, the response of ventral cells to Rho1 activation is similar to the response of wildtype cells in the dorsal epithelium (***Figure 4b v***. ***Figure 3d***-Activated Dorsal, ***Figure 3–Figure Supplement 4***). Thus, Dorsal is required to predispose ventral cells to constrict anisotropically along the anterior-posterior axis of the embryo.

**Figure 4.**
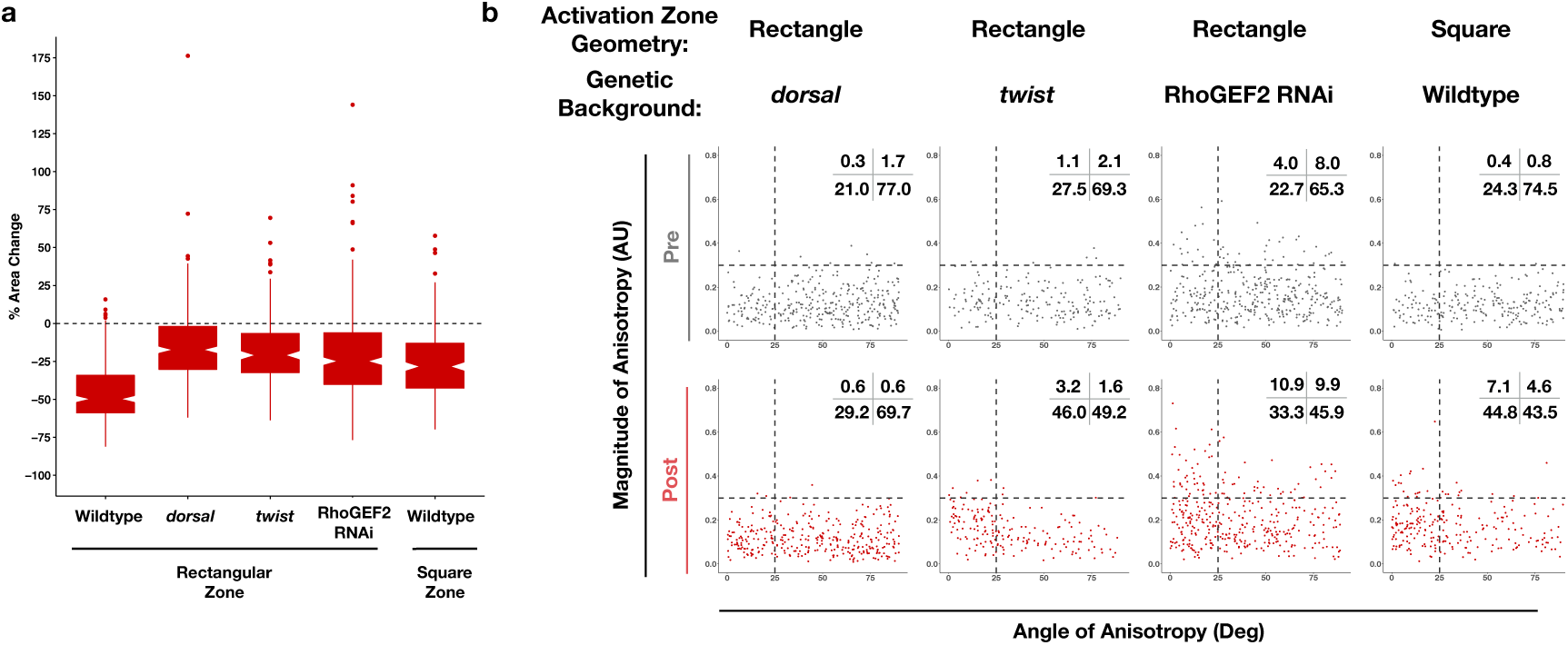
Dorsal is required for and Twist promotes aligned, anisotropic apical constriction in response to ectopic Rho1 activation. a) Quantification of apical area change induced by optogenetic Rho1 activation in wildtype and mutant backgrounds. Note: The wildtype, rectangular activation zone data is repeated from ***Figure 3c*** to facilitate comparison. b) Anisotropy scatter plots, as in ***Figure 3d***, for wildtype embryos subjected to a square region of ectopic Rho1 activation or specified mutant embryos subjected to a rectangular zone of activation. Insets show percentage of cells in each quadrant. Cells in the upper left quadrant exhibit highly aligned, anisotropic apical constriction. See ***Figure 3–Figure Supplement 4*** for plots of the changes in anisotropy of individual cells. 343 cells from 4 *dorsal* embryos, 189 cells from 3 *twist* embryos, 375 from 5 RhoGEF2 depleted embryos, and 239 cells from 5 square zone embryos were analyzed.

The transcription factor Twist is downstream of Dorsal activity in ventral cells. We optogenetically activated Rho1 in ventral cells of embryos homozygous for a null allele of *twist*. These mutant cells exhibited apical constriction following ectopic Rho1 activation (***Figure 4a***), but the amount of constriction, magnitude of anisotropy, and the degree of alignment with the anterior-posterior axis was less than that of wildtype ventral cells (***Figure 4b v***. ***Figure 3d***-Activated Ventral ; ***Figure 3–Figure Supplement 4***). Thus, Twist promotes the predisposition of ventral cells to constrict anisotropically along the anterior-posterior axis of the embryo. However, we note that ventral cells lacking Twist exhibit more aligned apical constriction than ventral cells lacking Dorsal (***Figure 4b***, ***Figure 3–Figure Supplement 4***). These distinct responses suggest that there is a Twist-independent mechanism downstream of Dorsal that promotes aligned anisotropic apical constriction in response to Rho1 activation. We speculate that Snail is responsible for this Twist-independent behavior, but repeated attempts to combine a null *snail* allele with our optogenetic components failed, so we were not able to test this hypothesis.

Dorsal is required for and Twist promotes ventral cells to respond to ectopic Rho1 activation with strong, aligned anisotropic apical constriction. This may reflect that the transcriptional targets of Dorsal and Twist are required for Rho1 activation during endogenous ventral furrow formation. Thus, we tested whether Dorsal and Twist are required for anisotropic apical constriction independent of their role in activating Rho1 by optogenetically activating Rho1 in ventral cells depleted of RhoGEF2, the endogenous activator of actomyosin contractility during ventral furrow formation. RhoGEF2 is required for proper organization of the actomyosin cytoskeleton during cellularization, and embryos lacking RhoGEF2 have some cellularization defects, contributing to irregularities in the epithelium (***Padash Barmchi et al., 2005***). Thus, a subset of cells depleted of RhoGEF2 are anisotropic, though randomly aligned, before optogenetic activation (***Figure 4b***). Despite the non-uniformity in these epithelia, optogenetic activation of Rho1 in ventral cells depleted of RhoGEF2 increased the extent of aligned, anisotropic apical constriction (***Figure 4, Figure 3–Figure Supplement 4***). Ectopic invaginations induced in the ventral or dorsal epithelium of embryos lacking RhoGEF2 failed to revert following cessation of optogenetic activation, in contrast to the rapid reversion of ectopic deformations in otherwise wildtype tissues. This suggests RhoGEF2 makes a significant contribution to the tension in the epithelium, likely through its role in organizing the actomyosin cytoskeleton. These results suggest ventral cells can respond to optogenetic Rho1 activation with anisotropic apical constriction in the absence of endogenous Rho1 activity. However, elevated Rho1 levels may contribute to strong, aligned, anisotropic apical constriction during endogenous ventral furrow formation.

Our experiments in the dorsal epithelium suggest that an asymmetric zone of Rho1 activation is not always sufficient to generate aligned, anisotropic apical constriction. However, we wondered whether an asymmetric zone of Rho1 activation might contribute to the cell shape changes seen in ventral epithelial cells during ventral furrow formation. To address this question, we locally activated Rho1 in a square region in ventral cells before any obvious apical constriction. Rectangular activation regions result in more highly anisotropic constrictions than square activation regions (***Figure 4b v***. ***Figure 3e***-VentralPost, ***Figure 3–Figure Supplement 4***). Thus, even though an asymmetric zone of Rho1 activation is not sufficient to induce anisotropic apical constriction in the dorsal epithelium, the asymmetry of the zone of Rho1 activation promotes the highly aligned anisotropic apical constriction in the ventral epithelium.

Taken together, these results suggest that both an asymmetric zone of Rho1 activation and ventral-specific factors, genetically downstream of Dorsal and Twist, contribute to the ability of ventral cells to respond to ectopic Rho1 activation with aligned, anisotropic apical constriction.

### Spreading of deformations within the endogenous ventral furrow region

We next asked whether optogenetic Rho1 activation would affect an already invaginating ventral furrow. Activation of Rho1 in a subset of cells locally accelerates their invagination (***Figure 5a***). This suggests Rho1 activity is rate-limiting during the invagination of the endogenous ventral furrow. Notably, the invagination of neighboring ventral furrow cells, outside of the defined activation region is also accelerated (***Figure 5a***-red arrow). Furthermore, optogenetic activation of Rho1 in the ventral epithelium prior to the onset of invagination frequently induces the invagination of both cells inside and neighboring the activation region (***Figure 5b***-red arrow). These non-autonomous cellular responses are not observed in the dorsal epithelium (***Figure 5c***). This ventral-specific response occurs in less than a minute, a time scale that is consistent with mechanical, rather than mechanochemical, transmission of forces. Thus, cells within the ventral and dorsal epithelia may exhibit differential mechanical properties.

**Figure 5.**
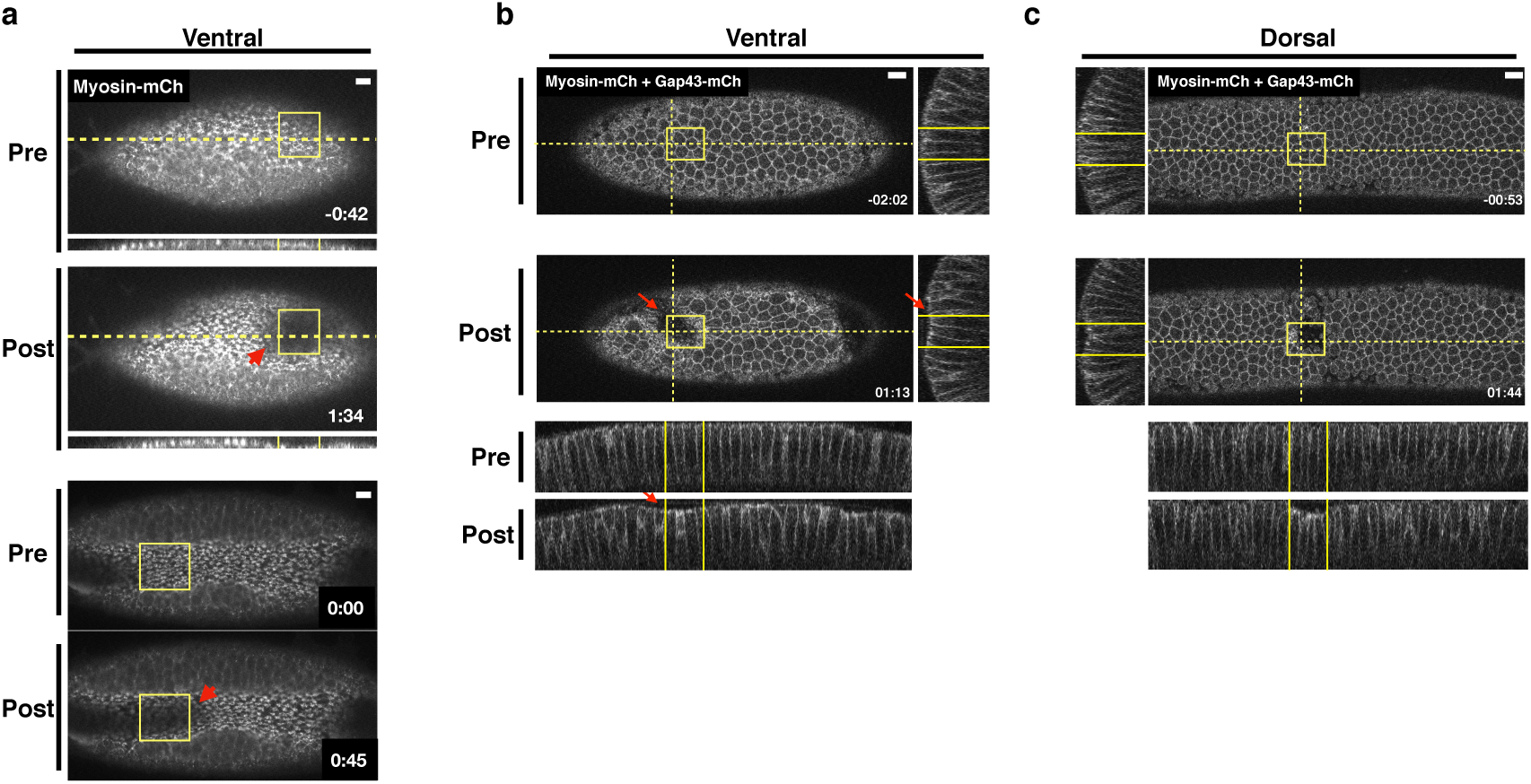
Optogenetic Rho1 activation specifically induces cell non-autonomous responses in the ventral epithelium. a) Ventral epithelium of an embryos expressing the optogenetic components and Myosin-Ch and exhibiting an established furrow. Ectopic Rho1 activity accelerates the invagination of the endogenous ventral furrow. This acceleration extends outside of the zone of Rho1 activation (red arrow). Data representative of 3/5 embryos. b-c) Ventral (b) and dorsal (c) epithelium of an embryo expressing LOVSsrA, PR-GEF, Gap43-mCh, and Myosin-Ch at the onset of gastrulation. Optogenetic activation of Rho1 within the yellow box induces an ectopic invagination. This ectopic invagination extends outside the defined activation region in the ventral (b, red arrow) but not dorsal (c) epithelium. Data representative of 5/8 ventral and 4/4 dorsal embryos. Scale bars are 10 μm. **Figure 5–video 1.** Movie of embryo shown in Figure 5a-bottom.

### Differential responses of cells flanking the Rho1 activation zone in the dorsal and ventral epithelium

Collectively, the results presented here suggest that ventral and dorsal cells exist in distinct mechanical environments. To further explore this possibility, we generated ectopic zones of Rho1 activation and focused on the behavior of cells adjacent to these zones. In the ventral epithelium, we observed extensive bending of non-activated cells towards optogenetically-induced invaginations (***Figure 6b***, filled arrowheads). This bending is readily visualized in maximum projections of the ventral surface post optogenetic activation, and it routinely extends several rows outside of the zone of photoactivation (***Figure 6b***, filled arrowheads). In contrast, long-range bending toward the ectopic invagination is not observed in the dorsal epithelium. Rather, the cells immediately adjacent to ectopic dorsal invaginations exhibit substantial stretching of their apical surfaces (***Figure 6a***, open arrowheads).

**Figure 6.**
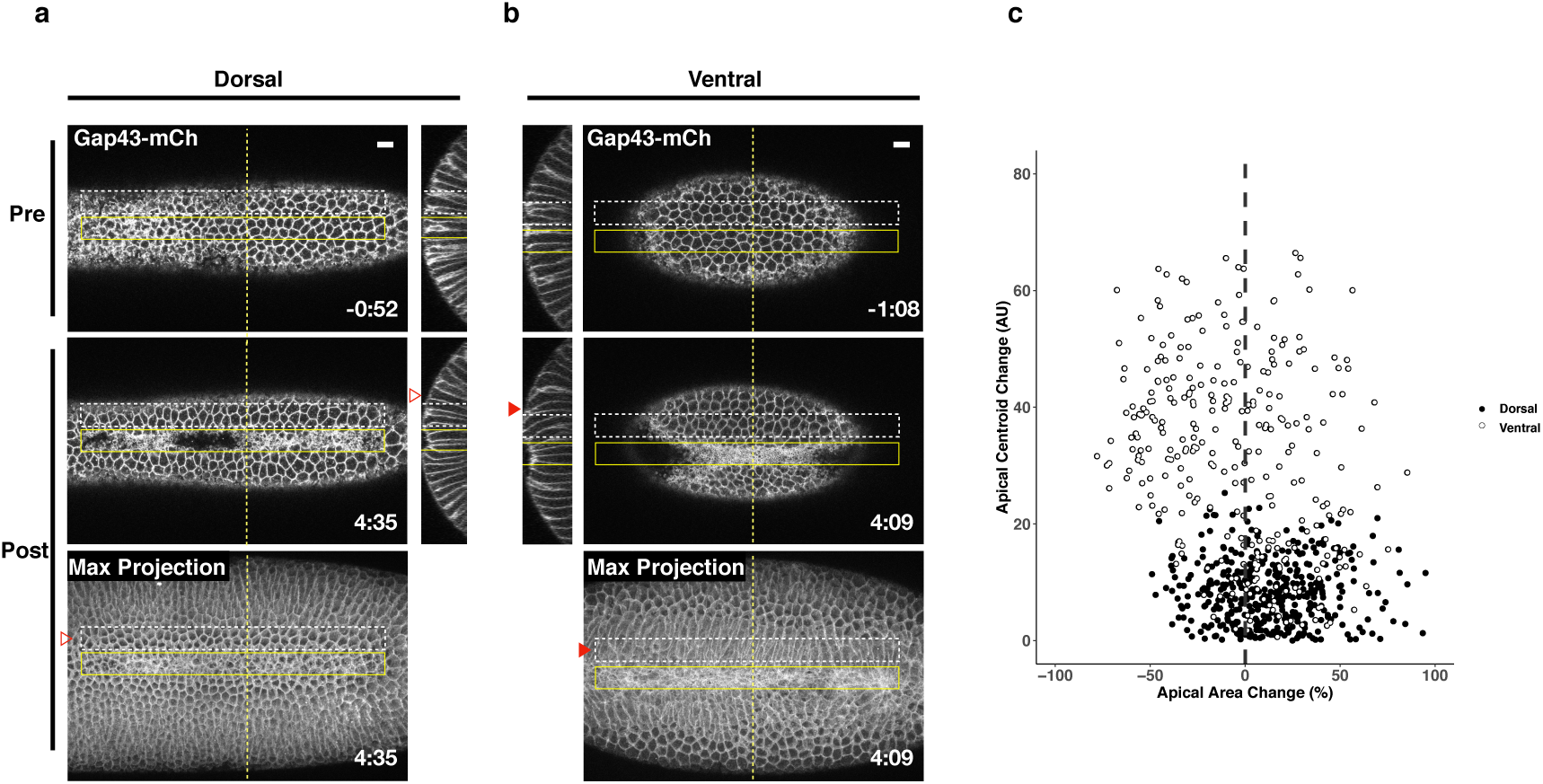
Non-activated cells bend towards ectopic invaginations specifically in the ventral epithelium. a-b) Dorsal (a) and ventral (b) epithelium of embryos expressing the optogenetic components and Gap43-mCh at the onset of gastrulation. Local Rho1 activation within the yellow box induces ectopic invaginations in both the dorsal and ventral epithelium. Bottom panels: Maximum intensity projections of the indicated time point. Data representative of 4/4 dorsal and 4/4 ventral embryos. c) Quantification of cell shape changes exhibited by cells neighboring ectopic invaginations. X axis is the percent apical area change; Y axis is the change in position of the centroid of the apical cell surface along the dorsal-ventral axis. 407 dorsal cells from 4 embryos and 298 ventral cells from 4 embryos were quantified. The white dashed boxes in a and b indicate the “neighbor” cells that were quantified for these two embryos. Filled arrowheads indicate long-range bending of ventral cells; open arrowheads indicate the corresponding cells in the dorsal epithelium. Dorsal cells at the periphery are not perpendicular to the imaging plane before or after photoactivation and therefore do not appear as hexagons in the maximum projection images. Time zero indicates the first pulse of blue light activation. Scale bars are 10 μm.

We quantified the bending of non-activated cells towards ectopic invaginations by measuring the change in the position of their apical centroids along the dorsal-ventral axis during the induction of the ectopic invaginations. Consistent with our visual observations, the centroids of the apical surface of non-activated ventral cells move substantially during the invagination of the photoactivated region, while the centroids of the apical surface of non-activated dorsal cells exhibit little movement during the comparable time (***Figure 6c***). Notably, dorsal cells neighboring ectopic invaginations are strongly biased towards expanding their apical surfaces, while the majority of ventral cells exhibit contraction of apical surfaces. Thus, not only do activated ventral and dorsal cells respond distinctly to optogenetic activation of Rho1, but the ventral and dorsal cells neighboring these regions of ectopic Rho1 activation respond distinctly to ectopic invaginations. These differential responses both within and adjacent to the activation zones suggest that the two epithelia exhibit differential mechanical properties.

## Discussion

Given the extensive evidence implicating Rho1 activation in ventral furrow formation, we assessed whether an asymmetric zone of Rho1 activity is sufficient to initiate this morphogenetic process in the *Drosophila* embryo. Optogenetic activation of Rho1 in the dorsal epithelium does not recapitulate all cell- and tissue-level aspects of ventral furrow formation. However, Rho1 activation in the ventral epithelium induces precocious furrowing that mirrors the endogenous process. We propose that this context-dependent response to ectopic Rho1 activation arises from distinct material properties of the dorsal and ventral epithelia.

### A robust, ubiquitously expressed optogenetic system for use in *Drosophila*

The LOV-domain based optogenetic probe generated in this study is expressed ubiquitously throughout the *Drosophila* lifecycle. This ubiquitous and non-perturbing expression allows Rho1 activation to be readily controlled in any *Drosophila* tissue without the need to combine the probe with tissue-specific drivers. This probe acts rapidly, inducing Rho1 activity within a minute of photoactivation. Precise spatial control of Rho1 activation can be induced using a range of standard fluorescent imaging methods. Ectopic deformations induced by optogenetic Rho1 activation in the dorsal embryonic epithelium are limited to the zone of optogenetic Rho1 activation, and, in the wing peripodial epithelium, Rho1 can be activated with subcellular precision.

### Optogenetically-induced invaginations are reversible

Using this optogenetic approach, we demonstrate that ectopic Rho1 activation is sufficient to induce ectopic, tissue-level shape changes throughout the embryonic epithelium at the onset of gastrulation. The cell shape changes induced by optogenetic Rho1 activation in ventral cells closely mirror those seen during endogenous ventral furrow formation, and ectopic Rho1 activation can modulate endogenous ventral furrow formation. This suggests that the potency of optogenetic activation of Rho1 via the LOV probe is on par with endogenous Rho1 activation during ventral furrowing.

Deformations induced by optogenetic activation of Rho1 persist through the duration of optogenetic activation. However, invaginated cells rapidly revert to their pre-activation positions and expand their apical areas following cessation of photoactivation, concurrent with rapid dissipation of optogenetically-induced myosin. Similar reversibilty occurs in the other tissues we examined as well as in cultured cells (***Wagner and Glotzer, 2016***; ***Oakes et al., 2017***). This reveals the existence of potent, widespread mechanisms for inactivating Rho1 and its effectors. We infer that ventral furrow formation is driven by sustained Rho1 activation that overcomes this global inhibition.

### PR-GEF and RhoGEF2-CRY2 induce distinct cellular responses

Our results are partially consistent with previous work, which activated Rho1 via membrane recruitment of a light-responsive RhoGEF2-CRY2 fusion protein (***Izquierdo et al., 2018***). Both optogenetic systems induce ectopic deformations in the dorsal embryonic epithelium, but only RhoGEF2-CRY2 induces pulsatile Rho1 activity and anisotropic apical constriction in the dorsal epithelium.

The two systems use different RhoA/Rho1-specific GEFs, and this may underlie the differing results; LOV recruits LARG(DH) while CRY2 is fused to RhoGEF2(DHPH). Despite LARG being an extremely potent RhoA activator *in vitro* (***Jaiswal et al., 2013***), the transgene expressing LARG(DH) is well tolerated (***Table 6***), suggesting this recruitable GEF is non-perturbing. To directly compare LARG and RhoGEF2, we generated flies expressing SspB-GFP-RhoGEF2(DHPH) from the same genomic location as PR-GEF. This transgene does not readily homozygose even in the absence of the LOVSsrA membrane anchor (***Table 6***), suggesting it has significant light-independent activity. PH domains of the GEF subfamily that includes RhoGEF2 and LARG bind RhoA-GTP, and, *in vitro*, the interaction between the PH domain and membrane-bound RhoA-GTP potentiates GEF activity by up to 40 fold (***Chen et al., 2010***; ***Medina et al., 2013***). Introducing two point mutations (F1044A, I1046E) into the PH domain of RhoGEF2, which are predicted to disrupt its binding to RhoA-GTP, allows the resultant transgene to readily homozygose (***Table 6***). These observations are consistent with RhoGEF2-CRY2 acting via a feedforward mechanism where it can be recruited by Rho1-GTP via its PH domain and thereby amplify Rho1-GTP. The ability of RhoGEF2-CRY2 to amplify both endogenous and light-induced Rho1 activity would be predicted to be particularly potent when it is overexpressed from a UAS promoter via Gal4. Feedforward activation via RhoGEF2-CRY2 may combine with the aforementioned mechanisms for Rho1 inactivation to generate the pulsatile Rho1 activity observed with RhoGEF2-CRY2 (***Izquierdo et al., 2018***). Amplification of endogenous Rho1 activity by RhoGEF2-CRY2 could also explain the anisotropic apical constrictions induced when this probe is optogenetically activated in the dorsal epithelium. Activated cells in this epithelium would need to deform against increased resistive forces exerted by their neighbors as a result of chronic Rho1 activation.

Although the GEF domain of RhoGEF2 is perturbing when over-expressed as an isolated domain, in the context of the full length protein, its ability to generate positive feedback via its PH domain may contribute to the Rho1 activity pulses observed during ventral furrow formation (***Martin et al., 2009***; ***Mason et al., 2016***).

### Requirements for ventral-specific responses to Rho1 activation

Despite the ability of asymmetric zones of Rho1 activation to induce deformations in both dorsal and ventral embryonic epithelia, they only induced strong, aligned, anisotropic apical constriction in the ventral epithelium. Dorsal is required for and Twist promotes this ventral-specific response, consistent with the idea that this property is a consequence of the gene expression differences that result from dorsal-ventral patterning. Twist is required to stabilize Rho1-driven apical constriction (***Martin et al., 2009***). Here, Twist promotes anisotropic apical constriction induced by sustained Rho1 activation. While it is possible that these two defects result from loss of the expression of a single Twist target gene, it is perhaps more likely that Twist controls the expression of multiple genes that independently contribute to ventral furrow formation. Notably, ventral cells lacking the Dorsal protein behave nearly identically to dorsal cells in wildtype embryos, while ventral cells lacking Twist exhibit weakly aligned, anisotropic apical constriction. Thus, a Twist-independent mechanism for generating aligned, anisotropic apical constriction must also exist. We speculate that Snail may also contribute to ventral-specific behavior.

Alternatively, Dorsal and/or Twist may be required for anisotropic apical constriction because each factor promotes Rho1 activation by RhoGEF2. However, ventral cells depleted of RhoGEF2 exhibit an increase in magnitude and alignment of anisotropy following ectopic Rho1 activation; thus, elevated Rho1 activity alone does not explain this ventral-specific response. The muted change in anisotropy of ventral cells lacking RhoGEF2 compared to wildtype ventral cells can be explained by the fact that cells depleted of RhoGEF2 exhibit higher degrees of anisotropy prior to optogenetic activation of Rho1, most likely because the epithelium is disorganized due to defects in cytoskeletal organization and cellularization (***Padash Barmchi et al., 2005***). Future work should identify the molecular targets of Dorsal and Twist that mediate anisotropic apical constriction. Two candidates of particular interest are Rap1 and its GEF Dzy; ventral cells lacking either of these proteins exhibit more isotropic apical constriction than wildtype ventral cells (***Sawyer et al., 2009***; ***Spahn et al., 2012***).

### Ventral and dorsal epithelia exhibit different material properties

The response of embryonic epithelial cells to optogenetic Rho1 activation depends on their location within the epithelium. Specifically, ventral, but not dorsal, cells constrict anisotropically, ventral deformations spread outside the activated regions, and several rows of epithelial cells bend toward activated ventral regions. Thus, the differences seen upon Rho1 activation are not limited to the response of the activated cells to Rho1 activation.

We propose that these ventral-specific behaviors arise as a consequence of dorsoventral patterning that endows the ventral epithelium with material properties that are distinct from those of the dorsal epithelium. These material properties (e.g. stiffness, deformability) likely result from differential organization and dynamics of the cytoskeleton and the junctions linking the cytoskeletons of neighboring cells. This dorsoventral patterning appears to specify the length scale over which forces are transmitted through the tissue. Importantly, these properties do not solely result from Rho1 activation in ventral cells, as RhoGEF2-depleted cells retain some ventral characteristics. We suggest that these material properties shape the reciprocal interactions between Rho1-activated cells and their neighbors, influencing the response both within and outside the Rho1 activated region.

The molecules responsible for ventral-specific material properties are not known, but it may include regulated cell-cell adhesion and the associated cytoskeletal networks. During ventral furrow formation, E-cadherin molecules in ventral cells reorganize from a sub-apical position to an apical position and become more densely packed (***Weng and Wieschaus, 2016***). These junctional rearrangements may contribute to efficient transmission of intracellular contractility throughout the ventral epithelium. Junctions transmit these forces through interactions with the actomyosin cytoskeleton which in turn influence the behavior of adherens junctions (***Weng and Wieschaus, 2016***). These interactions ultimately generate the supracellular actomyosin network observed during ventral furrow formation (***Martin et al., 2010***; ***Yevick et al., 2019***). In our experiments, ectopic Rho1 activation was not sufficient to induce such networks in the dorsal epithelium, indicating a requirement for ventral-specific factors in their assembly.

The cellular behaviors observed during light-induced invaginations are remarkably similar to those that occur during endogenous ventral furrowing (***Costa et al., 1994***; ***Leptin et al., 1992***; ***Leptin and Grunewald, 1990***; ***Sweeton et al., 1991***). These shape changes were widely thought to occur as a direct consequence of the transcriptional induction of Rho1-dependent contractility in the ventral epithelium. By comparing identical patterns and intensity of Rho1 activation in wildtype and mutant tissues, we have shown that dorsoventral patterning has additional relevant targets beyond Rho1 activation.

### Conclusion

In summary, this work shows that, despite inducing ectopic deformations, Rho1 activation alone is not sufficient to recapitulate the cell- and tissue-level behaviors observed during ventral furrow formation. Thus, a model of ventral furrow formation where Rho1 activity is the sole driver of cell and tissue behavior is incomplete. We propose that ventral-specific behaviors may arise from expression of factors that modulate the cytoskeleton and its connection to adherens junctions as well as promote strong intercellular coupling among cells of the ventral epithelium.

## Methods and Materials

### Plasmids

Plasmids used in this studied are listed in ***Table 1***. pUbi-stop-mCD8GFP containing an attB site and pUbi>mEGFP-Anillin(RBD) were gifts from T. Lecuit. Plasmids created for this study were generated using SLiCE (***Zhang et al., 2012***) or one-step isothermal *in vitro* recombination (***Gibson et al., 2009***). Stargazin-GFP-LOVpep and PDZx2-mCherry-LARG(DH) plasmids were published previously (***Wagner and Glotzer, 2016***). Venus-iLID-CAAX and tgRFPt-SspB WT were obtained from Addgene (60411, 60415). pMT>Gal4 (***Klueg et al., 2002***) was obtained from the Drosophila Genomics Resource Center.

**Table 1.** Plasmids Used

| Plasmid Generated | Fragment Source | Backbone Source |
| --- | --- | --- |
| pMT>Stargazin-GFP*-LOVSsrA<br>(* represents inactivation of fluorophore with Y66C mutation.) | Stargazin-GFP amplified from Stargazin-GFP-LOVpep<br>(*Wagner and Glotzer, 2016*)<br>LOVSsrA amplified from Venus-ILID-CAAX<br>(*Guntas et al., 2015*) (Addgene: 60411)<br>GFP silenced by<br>site-directed mutagenesis (Y66C) | pMT>Gal4 (*Klueg et al., 2002*) |
| pMT>Stargazin-GFP*-LOV(I427V)SsrA | N/A | pMT>Stargazin-GFP*-LOVSsrA<br>w/ site directed mutagenesis |
| pUbi>Stargazin-GFP*-LOVSsrA | Stargazin-GFP*-LOVSsrA amplified from<br>pMT>Stargazin-GFP*-LOVSsrA | pUbi-stop-mCD8GFP<br>(Contains attB site) |
| pUbi>Stargazin-GFP*-LOV(I427V)SsrA | Stargazin-GFP*-LOV(I427V)SsrA amplified from<br>pMT>Stargazin-GFP*-LOV(I427V)SsrA | pUbi-stop-mCD8GFP<br>(Contains attB site) |
| pUbi>SspB-GFP-LARG(DH) | LARG(DH) amplified from PDZx2-mCherry-LARG(DH)<br>(*Wagner and Glotzer, 2016*)<br>SspB amplified from tgRFPt-SspB(WT)<br>(*Guntas et al., 2015*) (Addgene: 60415) | pUbi-stop-mCD8GFP<br>(Contains attB site) |
| pMT>tagRFP-SspB | SspB amplified from tgRFPt-SspB(WT)<br>(*Guntas et al., 2015*) (Addgene: 60415) | pMT>Gal4 |
| pUbi>SspB-mScar | SspB amplified from<br>pMT>tagRFP-SspB<br>mScar amplified from<br>pmScarlet-C1 (Addgene: 85042) | pUbi-stop-mCD8GFP<br>(Contains attB site) |
| pUbi>SspB-GFP-RhoGEF2(DHPH) | RhoGEF2(DHPH) amplified from genomic prep of<br>Sp/CyO; UASp>RFP-RhoGEF2/TM3<br>(*Wenzl et al., 2010*) | pUbi-SspB-GFP-LARG(DH)<br>(Replace LARG(DH))<br>(Contains attB site) |
| pUbi>SspB-GFP-RhoGEF2(DHPH-F1044A,I1046E) | N/A | pUbi>SspB-GFP-RhoGEF2(DHPH)<br>w/ site directed mutagenesis |
| pUbi>mCherry-Anillin(RBD) | Anillin(RBD) amplified from<br>pUbi>mEGFP-Anillin(RBD) (*Munjal et al., 2015*)<br>mCherry amplified from<br>pm-Cherry2B (F. Valbuena and B. Glick,<br>manuscript in preparation) | pUbi-stop-mCD8GFP<br>(Contains attB site) |

### Fly stocks

*Drosophila melanogaster* was cultured using standard techniques at 25°C. Both male and female animals were used. Stocks used in this study include *pUbi>Gap43-mCherry/TM3*, generated by P-element insertion and was a gift from A. Martin; *pSqh>Sqh-mCherry* (***Martin et al., 2009***); Δ *halo AJ twist*^*EY* 53*R*12^/CyO, a gift from M. Leptin; *dl*^1^ *cn*^1^ *sca*^1^/CyO (BID: 3236); *UAS>RhoGEF2 shRNA* (BID: 76255); *P(mat-tub-Gal4)mat67* (BID: 7062).

Transgenic flies were generated by PhiC31-directed integration (GenetiVision). Transgenic lines generated for this study include: *Ubi>Stargazin-GFP*-LOVSsrA (attP2), Ubi>Stargazin-GFP*-LOV(I427V)SsrA (attP2), Ubi>SspB-GFP-LARG(DH) (VK37), Ubi>SspB-GFP-LARG(DH) (VK31), Ubi>SspB-GFP-RhoGEF2(DHPH) (VK37), Ubi>SspB-GFP-RhoGEF2(DHPH-F1044A, I1046E) (VK37), Ubi>SspB-mScarlet (VK37), Ubi>mCherry-Anillin(RBD) (attP40)*.

Genotypes of flies used in each experiment are listed in ***Table 2*** and ***Table 3***.

**Table 2.**
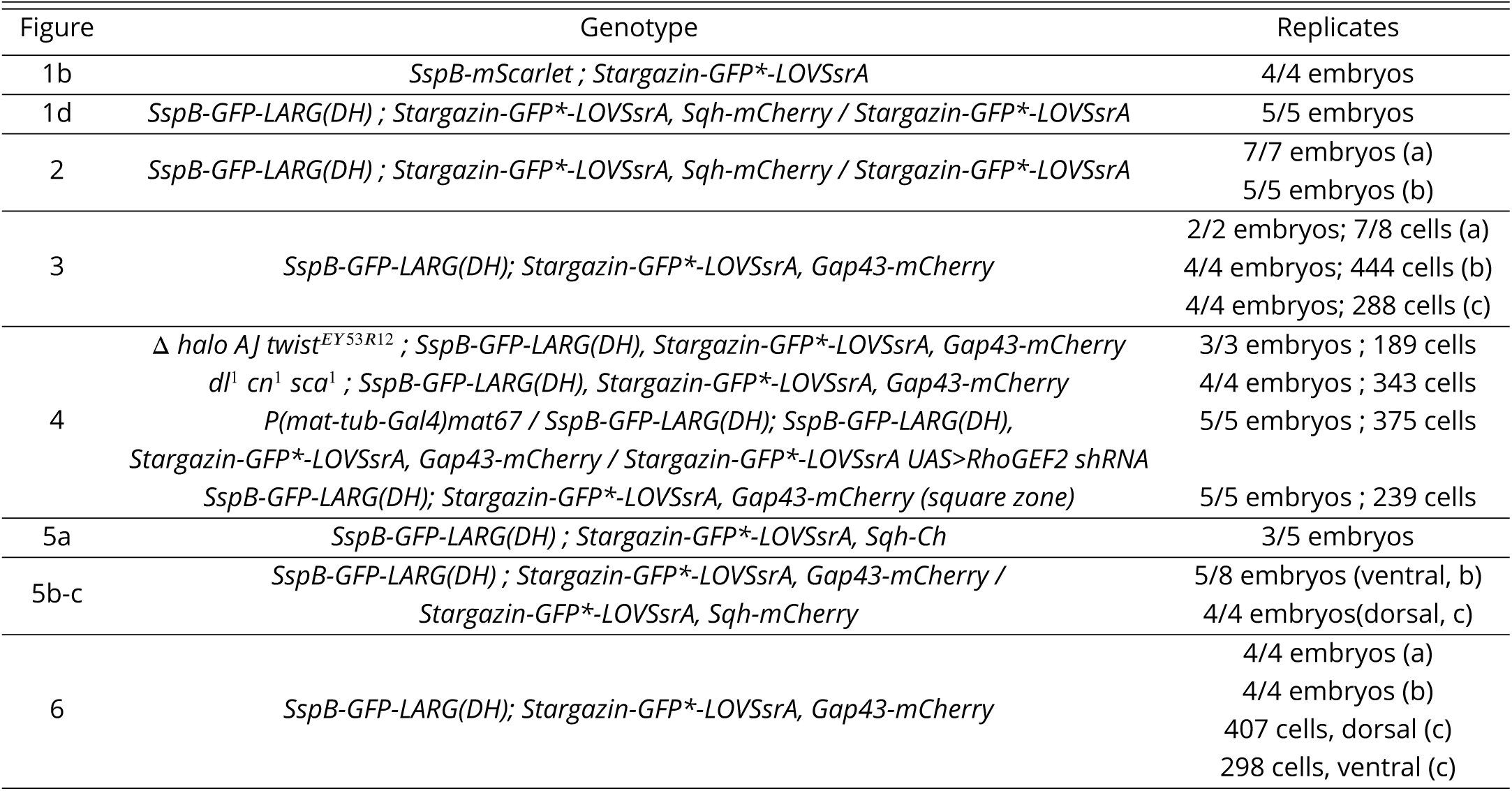
Genotypes and Reproducibility: Main Figures

**Table 3.**
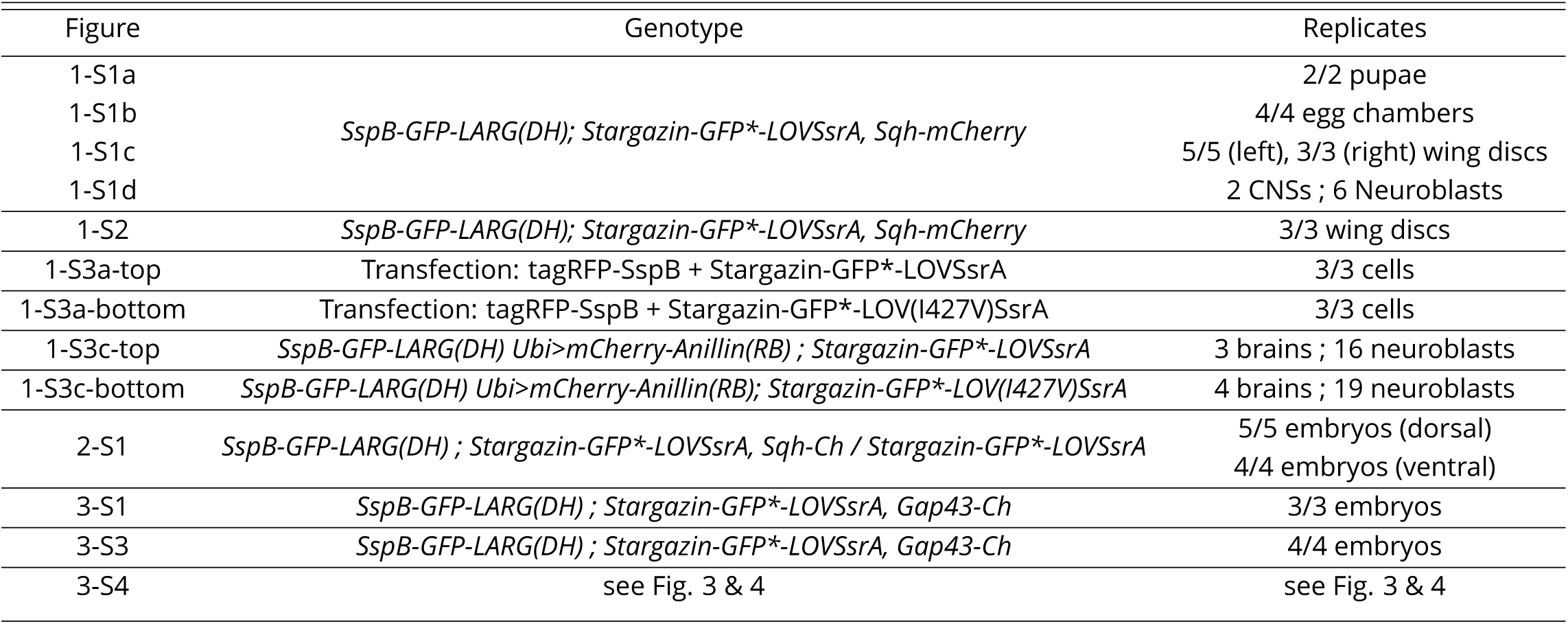
Genotypes and Reproducibility: Supplemental Figures

### S2 cells

3.1 x 10^6^ S2 cells were transfected with 100 ng pMT>tagRFP-SspB and 250 ng pMT>Stargazin-GFP*-LOVSsrA or 250ng pMT>Stargazin-GFP*-LOV(I427V)SsrA using dimethyldioctadecyl-ammonium bromide (Sigma) (***Han, 1996***) at 250 ug/mL in six well plates. Expression from the pMT promoter was induced 2 days after transfection by addition of 0.35 mM *CuSO*_*4*_. Cells were imaged live 24 hrs after *CuSO*_*4*_ induction. 50 μL of the S2 cell culture was plated on a glass slide and covered with a coverslip. Clay feet were used as spacers between the slide and coverslip. See ***Table 5*** for activation protocol details.

### Preparation of *Drosophila* tissues for live imaging

*Drosophila* embryos were collected on apple juice agar plates for 90 min and aged for 90-120 min at 25°C such that a majority of embryos were completing cellularization at the time of mounting. Embryos were dechorionated in 30% bleach for 1 min, rinsed in water, aligned on an apple juice agar pad, and mounted on a coverslip with embryo glue (adhesive from double sided tape dissolved in heptane). The imaged surface (dorsal or ventral) was mounted on the coverslip. This coverslip was affixed via petroleum jelly to a metal slide with a hole in the center. Embryos were covered with halocarbon oil 200 immediately after mounting; they were not compressed.

Central nervous systems were dissected from wandering third instar larvae in Schneider’s *Drosophila* Medium (Sigma) supplemented with 10% Fetal Bovine Serum (Thermo Fisher Scientific). Central nervous systems were imaged in a chamber comprising a coverslip affixed with petroleum jelly to a metal slide with a hole in the center. Following dissection, central nervous systems were mounted in the chamber such that their dorsal side contacted the coverslip. The chamber was flooded with Chan and Gerhing’s balanced solution (***Chan and Gehring, 1971***) to completely cover the central nervous system, and a gas-permeable membrane (YSI: 5793) was placed over the chamber to limit evaporation. These chambers were imaged on an inverted microscope.

Wing imaginal discs were dissected from wandering third instar larvae in S2 cell media supplemented with 10% FBS. Wing discs were mounted between a slide and glass coverslip in 50uL Chan and Gehring’s balanced solution. Clay feet were used as spacers between the slide and coverslip.

To prepare pupal nota, whole pupae were extracted from their pupal cases 18 hours post pupariation and mounted on a glass slide in a humid chamber, as described previously (***Zitserman and Roegiers, 2011***). Pupal nota were imaged on an upright microscope.

To image egg chambers, ovaries were dissected from 3-5 day old females aged on yeast. Individual stage 10 egg chambers were isolated and mounted between a coverslip and a slide. Clay feet were used as a spacer between the slide and coverslip.

### Live imaging and optogenetic experiments

Global activation experiments were performed on a 63x/1.4 numerical aperture (NA) oil immersion lens on a Zeiss Axiovert 200M equipped with a Yokogawa CSU-10 spinning disk unit (McBain) and illuminated with 50-mW, 473-nm and 20-mW, 561-nm lasers (Cobolt) or on a Zeiss Axioimager M1 equipped with a Yokogawa CSU-X1 spinning disk unit (Solamere) and illuminated with 50-mW, 488-nm and 50-mW, 561-nm lasers (Coherent). Images were captured on a Cascade 1K electron microscope (EM) CCD camera, a Cascade 512BT (Photometrics), or a Prime 95B (Photometrics) controlled by MetaMorph (Molecular Devices). Photoactivation was accomplished by illuminating the sample with 488 nm light for the indicated exposure times (***Table 4*** & ***Table 5***).

**Table 4.** Activation Protocols: Main Figures

| Figure | Microscope, Wavelength | Activation Protocol | Total Activation Time |
| --- | --- | --- | --- |
| 1b | LSM880, 488nm | Every 20 sec | 1 min 40 sec |
| 1d | Spinning Disc, 488nm | Global (1000ms) every 20 sec | 1 min |
| 2 a-b | LSM880, 488nm | Every 20 sec | 4min |
| 3a-b | LSM880, 488nm | Every 20 sec | 4 min |
| 4 | LSM880, 488nm | Every 20 sec | 4 min |
| 5a | LSM880, 488nm | Every 20 sec | 1 min 20 sec |
| 5b | LSM880, 488nm | Every 20 sec | 1 min 20 sec |
| 5c | LSM880, 488nm | Every 20 sec | 1 min 40 sec |
| 6a-b | LSM880, 488nm | Every 20 sec | 4 min |

**Table 5.**
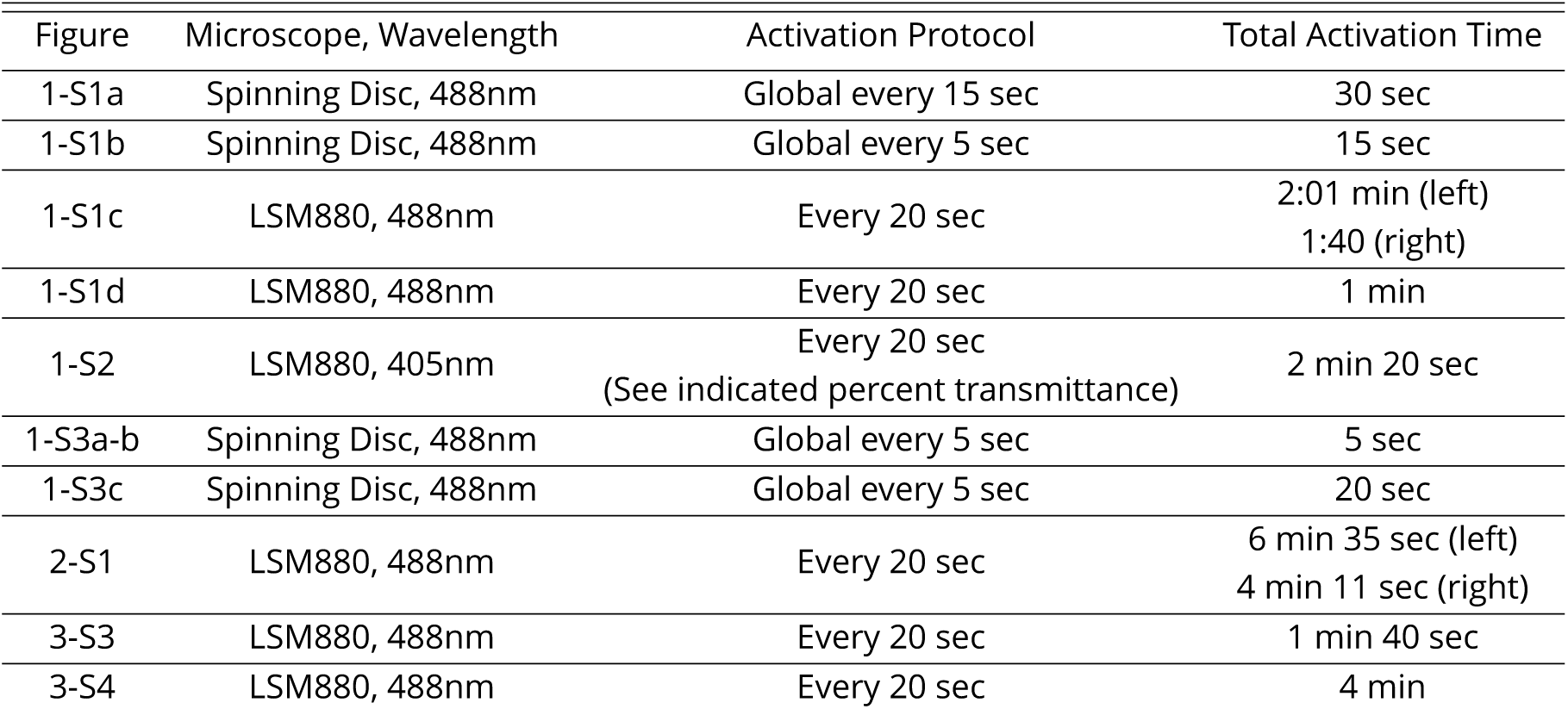
Activation Protocols: Supplemental Figures

**Table 6.** Recruitable GEF Viability Tests

| Cross Scored | Possible Genotype | Observed | Expected | Chi<sup>2</sup> |
| --- | --- | --- | --- | --- |
| SspB-GFP-LARG(DH)/CyO x | 1) LARG (DH)/CyO | 379 | 362.7 | 2.198 |
| SspB-GFP-LARG(DH)/CyO | 2) LARG(DH)/LARG(DH) | 165 | 181.3 |  |
| SspB-GFP-RhoGEF2(DHPH)/CyO x | 1) RhoGEF2(DHPH)/CyO | 318 | 248 | 59.27 |
| SspB-GFP-RhoGEF2(DHPH)/CyO | 2) RhoGEF2(DHPH)/RhoGEF2(DHPH) | 54 | 124 |  |
| SspB-GFP-RhoGEF2(DHPH-F1044A,I1046E)/CyO x | 1) RhoGEF2(DHPH*)/CyO | 430 | 414.7 | 1.694 |
| SspB-GFP-RhoGEF2(DHPH-F1044A,I1046E)/CyO | 2) RhoGEF2(DHPH*)/RhoGEF2(DHPH*) | 192 | 207.3 |  |

Local activation experiments were performed on a inverted Zeiss LSM880 laser scanning confocal microscope with a 40X/1.4 numerical aperture (NA) objective. mCherry or mScarlet fluorescence was excited using the 561 nM solid state laser and was detected via a GaAsP spectral detector. Activation regions, indicated with yellow boxes throughout this manuscript, were defined in the “Bleaching” module. Pixels within the defined activation zone were exposed to 488nm light attenuated to 0.01 or 0.1 percent laser transmittance, using an Acousto-optic tunable filter, for 15 iterations every 20 seconds for the duration of the activation period. In general, we acquired a “pre” Z-Series of Gap43-mCh or Sqh-mCh, activated the defined region with 488nm light in a single Z-plane, and acquired a “post” Z-Series of Gap43-mCh or Sqh-mCh. See ***Table 4*** & ***Table 5*** for specific activation protocols for each experiment.

### Image processing and cell shape analysis

All images were processed with FIJI (***Schindelin et al., 2012***). TissueAnalyzer (***Aigouy et al., 2010***), a FIJI plugin, was used to segment the embryonic epithelium and track cells for quantification of apical area, apical cell anisotropy, and apical cell centroid. “Pre” and “Post” Z-stacks were tracked separately in TissueAnalyzer, and data for the apical area, apical cell elongation (a proxy for anisotropy), and apical cell centroid were extracted from each timepoint and concatenated into a master database. Percent area change of the apical cell surface was calculated as (EndArea-StartArea)/StartArea*100. Magnitude of anisotropy, calculated in TissueAnalyzer, is a value ranging between 0 and 1, with 0 being highly isotropic and 1 being highly anisotropic. We converted the orientation of this anisotropy, calculated in TissueAnalyzer, to degrees for plotting (***Aigouy et al., 2010***). Data were plotted in RStudio with ggplot2.

## Acknowledgements

This work was supported by R01GM085087, R35GM12709, and a France and Chicago Collaborating through the Sciences grant (M.G.), R01NS034783 (R.G.F), and NIH T32 GM007183 and NSF GRFP DGE-1144082; DGE-1746045 (A.R). We thank Ed Munro for helpful comments on this manuscript. We thank Ed Munro, Sally Horne-Badovinac, Thomas Lecuit, M.G. lab members, and R.G.F. lab members for helpful discussions and support. Benoit Aigouy provided assistance with TissueAnalyzer. Audrey Williams helped with oocyte dissections. We thank the Glick, Martin, Leptin, and Lecuit labs for generous sharing of reagents. We thank Ben Glick for access to the SnapGene molecular biology software (http://www.snapgene.com). Stocks obtained from the Bloomington *Drosophila* Stock Center (NIH P40OD018537) were used in this study. Reagents obtained from *Drosophila* Genomics Resource Center, supported by NIH grant 2P40OD010949 were used in this study.

---

**Figure 1–Figure supplement 1.**
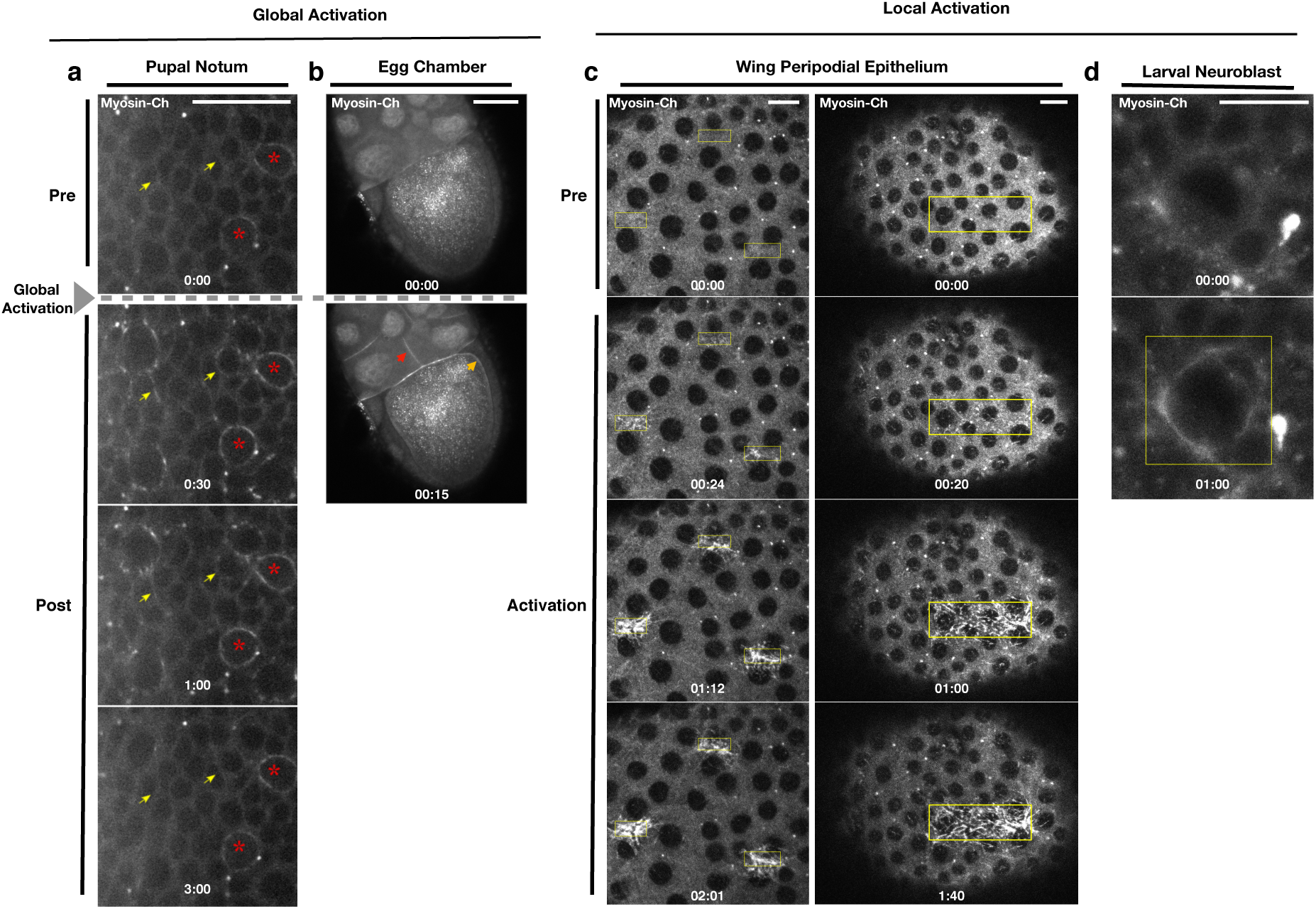
Recruitment of PR-GEF activates Rho1 in all tissues tested. a) Pupal Notum expressing the optogenetic probes. Myosin-Ch is shown before and after global photoactivation. Yellow arrow indicates the same cell junction over time. Asterisks indicate mitotic cells. Data representative of 2/2 pupae. b) Egg chambers expressing the optogenetic probes. Myosin-Ch is shown before and after global photoactivation. Nurse cell junctions (red arrow) and the oocyte cortex (yellow arrow) are indicated. Data representative of 4/4 egg chambers. c-d) Larval wing imaginal discs (c) and larval neuroblast (d) expressing the optogenetic probes. Rho1 was locally photoactivated within the yellow boxes. Myosin-Ch is shown before and during activation. Myosin-Ch accumulates with sub-cellular precision in the peripodial epithelium, consisting of squamous cells (c, left). Data representative of 5/5 wing discs (c, left), 3/3 wing discs (c, right), and 6/6 neuroblasts from 2 central nervous systems (d). Time zero indicates the first pulse of blue light activation. Scale bars are 10 μm.

**Figure 1–Figure supplement 2.**
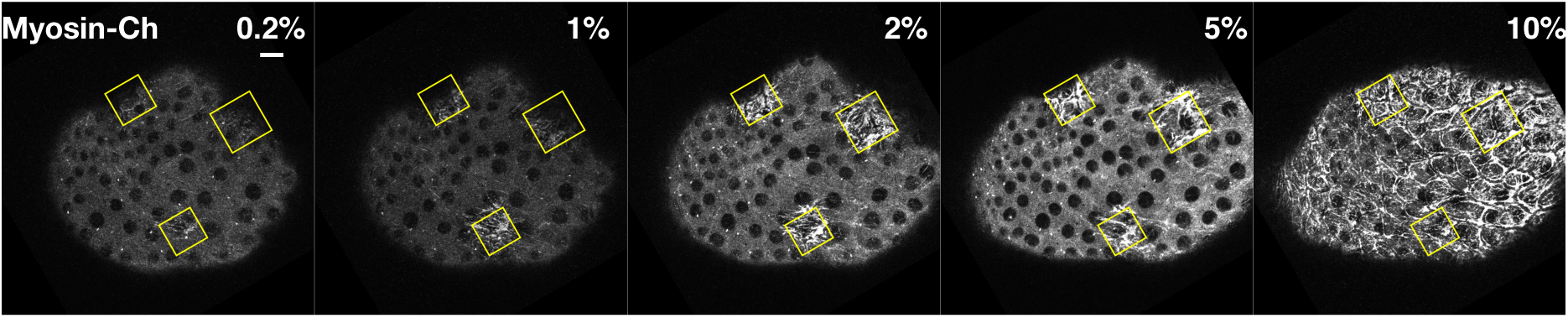
Ectopic Rho1 activation is sensitive to light dose. Larval wing peripodial epithelia expressing the optogenetic components and Myosin-Ch. Rho1 was optogenetically activated in the yellow boxes. Laser power was attenuated to the indicated percent transmittance using an acusto-optical tunable filter. 10% transmittance induces substantial Rho1 activation outside of the activation zone. Lowering the laser transmittance yields decreasing amounts of myosin accumulation. Photoactivation lasted 2 min 20 sec for each % transmittance. Data representative of 4/4 wing imaginal discs. Scale bars are 10 μm.

**Figure 1–Figure supplement 3.**
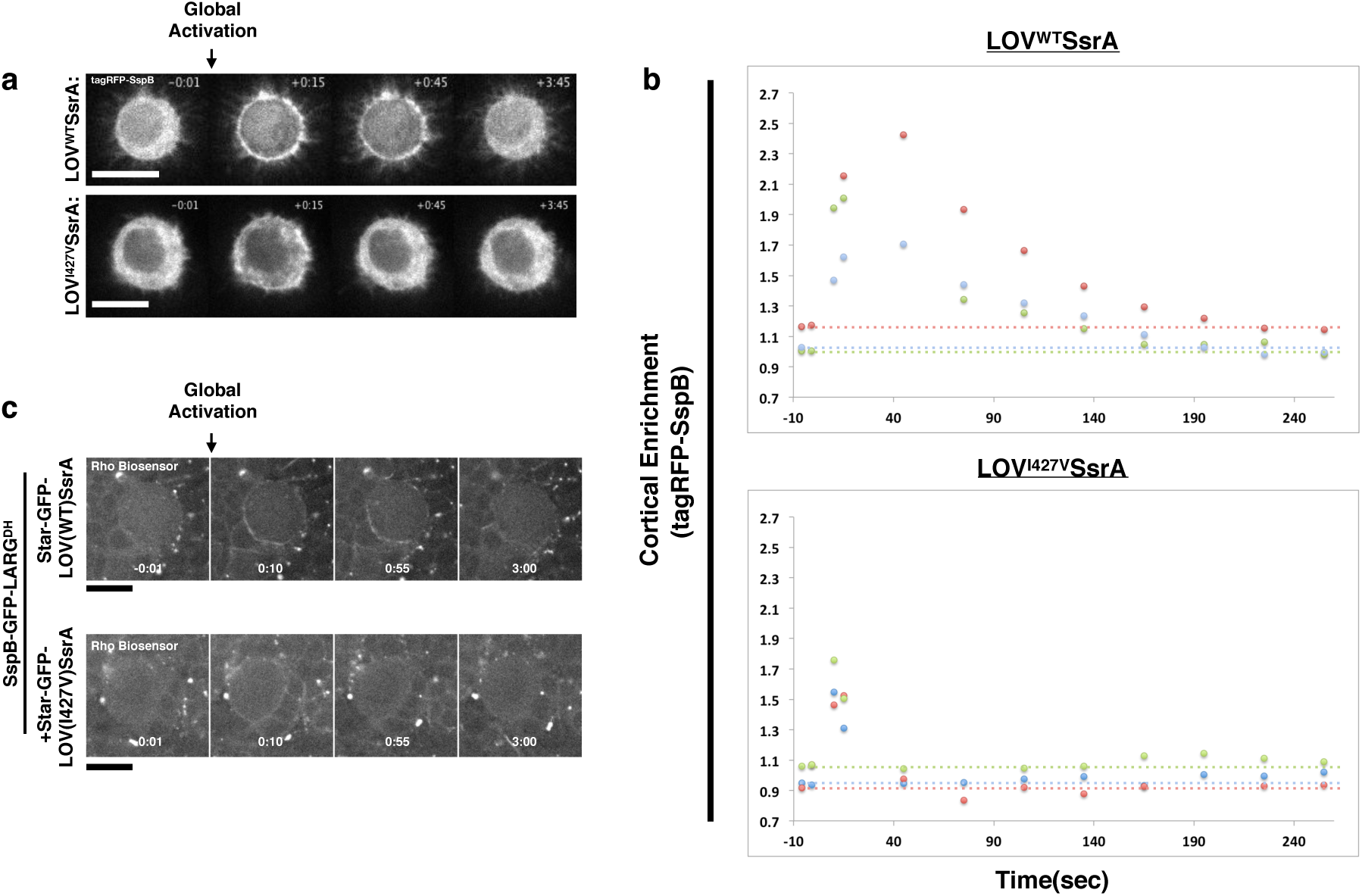
Inactivation kinetics of the LOV domain dictate the off rate of optogenetic-induced Rho1 activity. a) S2 cells transiently transfected with recruitable tagRFP-SspB and a membrane localized WT LOV domain (top) or fast-cycling (I427V) LOV domain (bottom). Representative cells are shown before and after global photoactivation. b) Quantification of the cortical enrichment (membrane/cytoplasm) of SspB-tagRFP following global optogenetic Rho1 activation. c) Larval neuroblasts expressing PR-GEF, Rho-biosensor, and WT (top) or fast-cycling (bottom) membrane-localized LOV domain shown before and after global photoactivation. Scale bars are 5 μm. Rho-biosensor consists of the Rho binding domain of Anillin, a RhoA effector, fused to mCherry (***Munjal et al., 2015***; ***Piekny and Glotzer, 2008***). Data representative of 16 neuroblasts from 3 brains (top) and 19 neuroblasts from 4 brains (bottom).

**Figure 2–Figure supplement 1.**
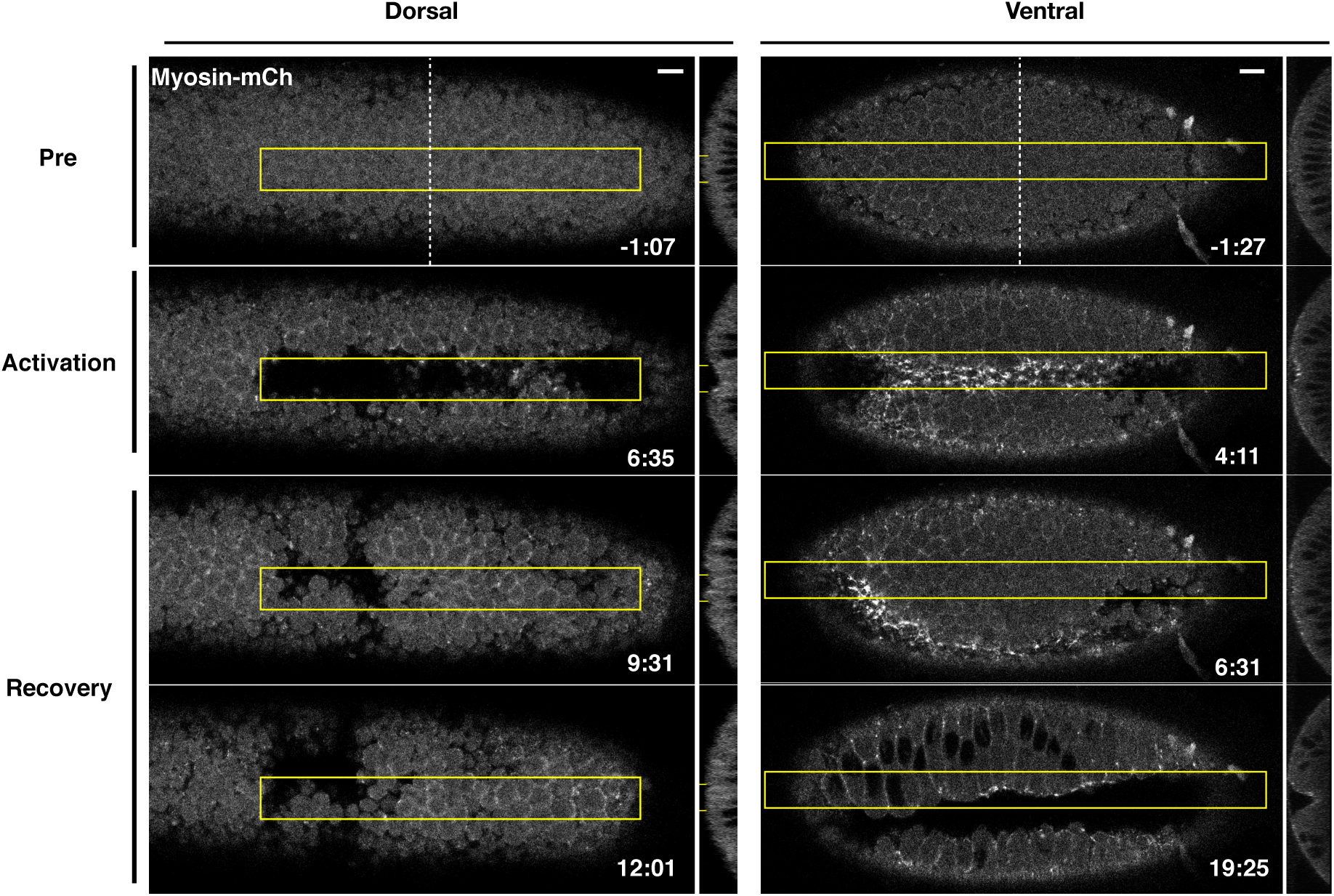
Optogenetic-induced invaginations revert following cessation of Rho1 activation. Dorsal (left) or ventral (right) epithelium of *Drosophila* embryos expressing the optogenetic components and Myosin-Ch before, during, and after local Rho1 activation within the yellow boxes. Note that some cells in the dorsal epithelium remain invaginated after the recovery period. These sustained pockets of invagination are sometimes seen where the dorsal transverse folds form. Data representative of 5/5 dorsal and 4/4 ventral embryos. Time zero indicates the first pulse of blue light activation. Scale bars are 10 μm.

**Figure 3–Figure supplement 1.**
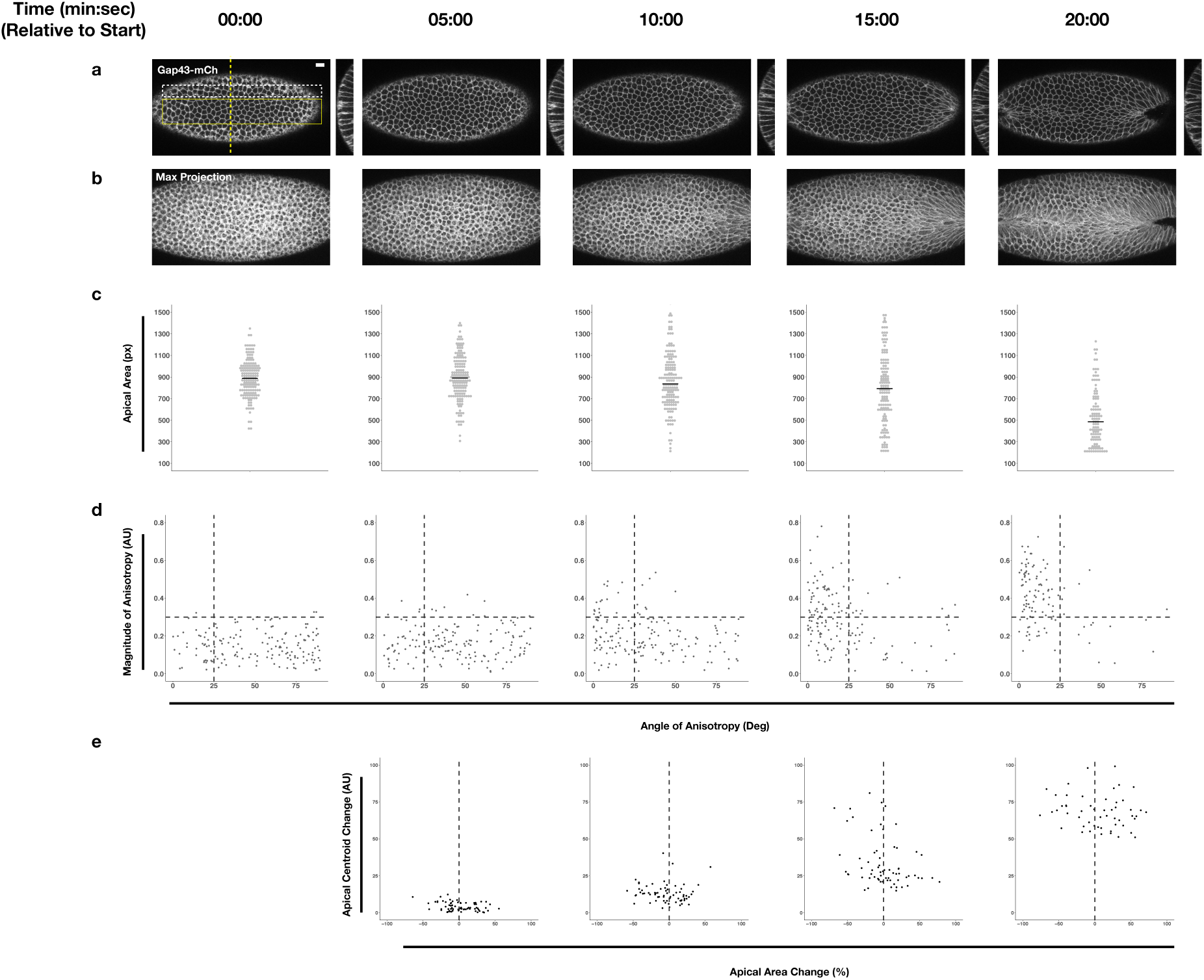
Quantification of endogenous ventral furrow formation. a-b) Single plane (a) and maximum projection (b) images of a non-activated embryo expressing the optogenetic components. c-d) Plot of apical area (c) or anisotropy (d) of cells in the endogenous ventral furrow (yellow box in a) at indicated time points. Black lines in (c) represent median. e) Plot of centroid change, relative to time 00:00, along the Y axis for cells neighboring the ventral furrow (white box in a). Data representative of 3/3 embryos. Scale bars are 10 μm.

**Figure 3–Figure supplement 2.**
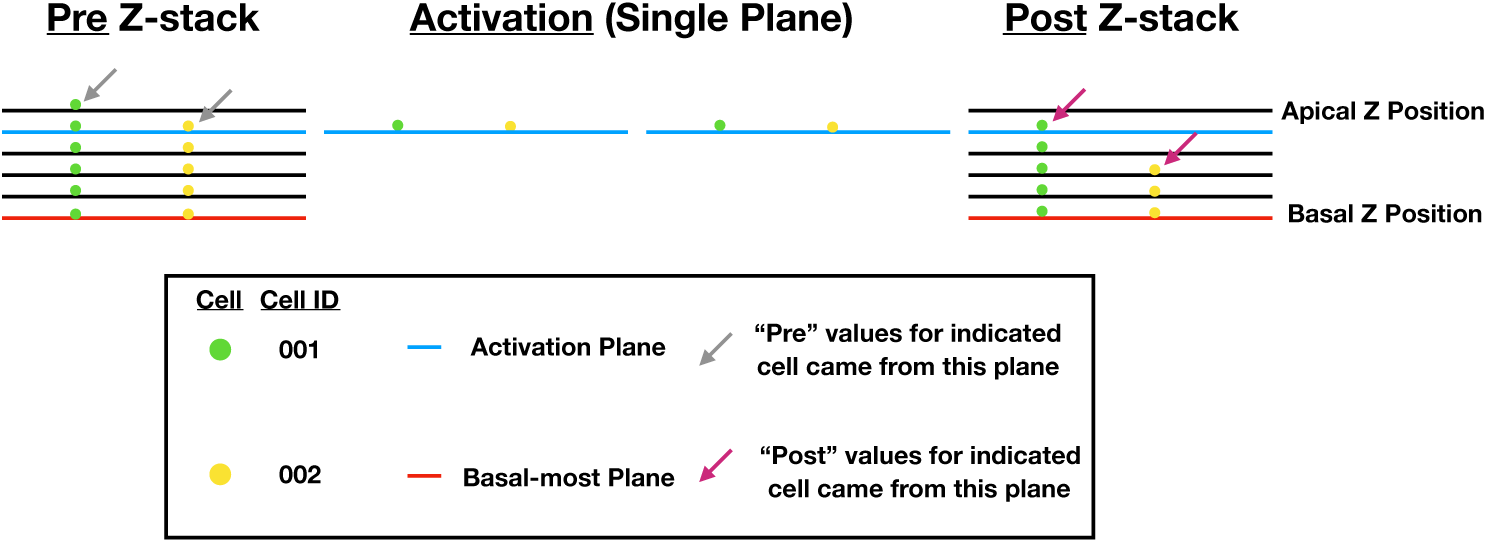
Schematic of data collection and analysis for local activation experiments.

**Figure 3–Figure supplement 3.**
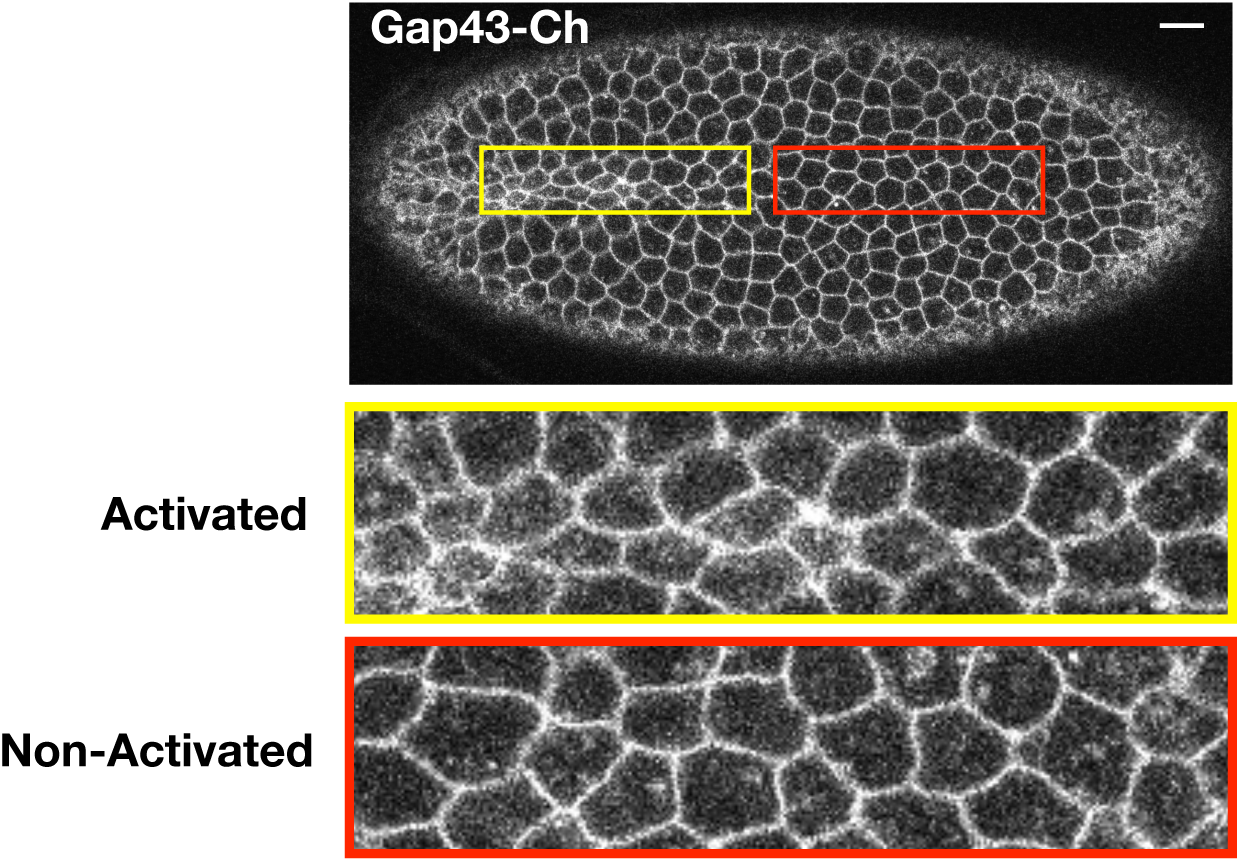
Optogenetic activation of Rho1 induces precocious cell shape changes in the ventral epithelium. Ventral surface of an embryo expressing the optogenetic components and Gap43-Ch. Rho1 was activated within the yellow box. Zoomed images of activated (yellow) and non-activated (red) cells are shown. Data representative of 4/4 embryos. Scale bars are 10 μm.

**Figure 3–Figure supplement 4.**
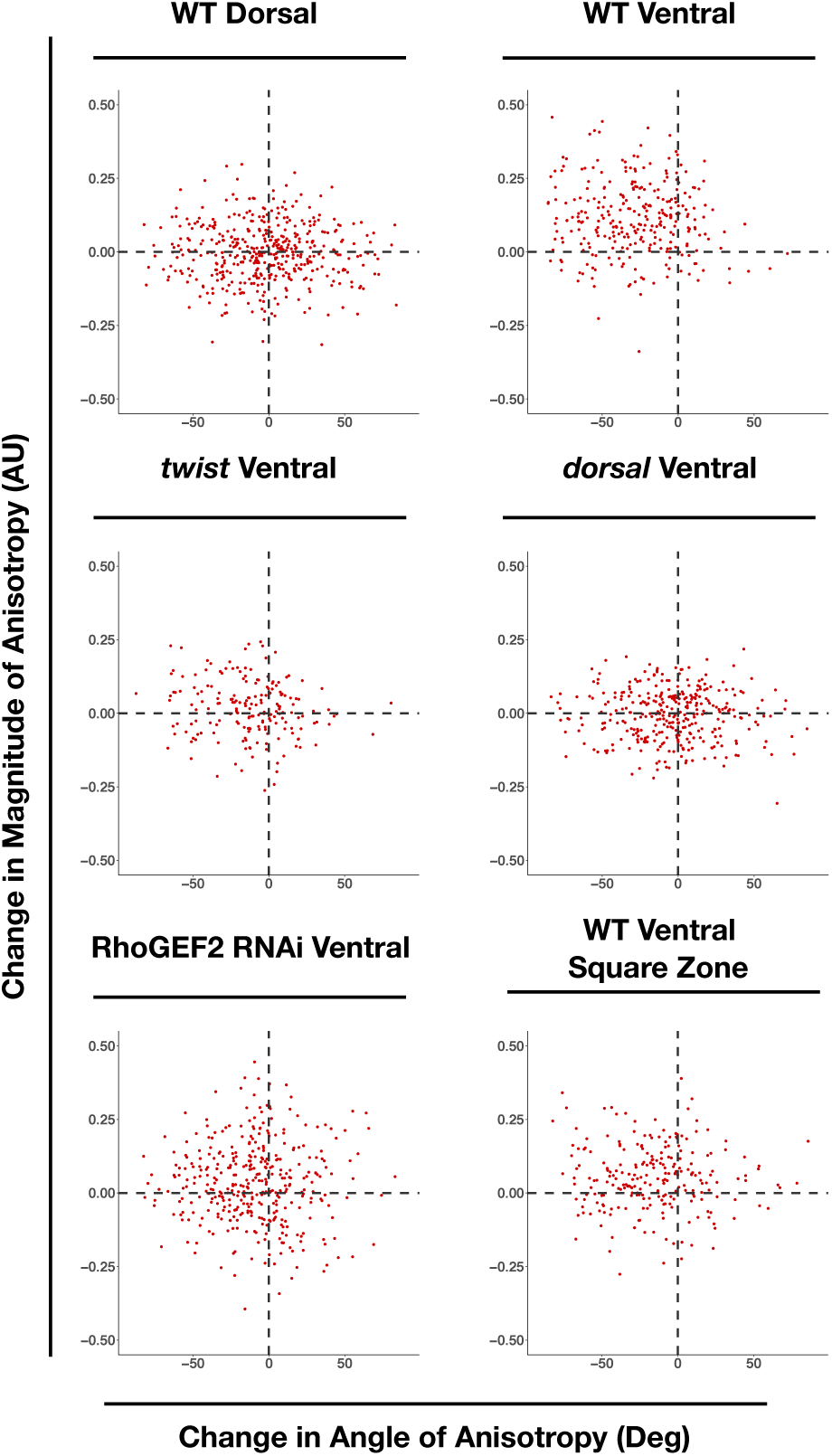
WT ventral cells exhibit large changes in the magnitude and alignment of anisotropy in response to Rho1 activation. X and Y axes show the changes in anisotropy angle (X) and magnitude (Y) for each cell over the course of optogenetic activation. Angle change was calculated as end angle (in degrees) minus start angle (in degrees). Magnitude change was calculated as end magnitude minus start magnitude. 444 cells from 4 wildtype dorsally oriented, 288 cells from 4 wildtype ventrally oriented, 343 cells from 4 *dorsal* embryos, 189 cells from 3 *twist* embryos, 375 cells from 5 RhoGEF2 depleted embryos, and 239 cells from 5 square zone embryos were analyzed.

## References

Aigouy, B., Farhadifar, R., Staple, D. B., Sagner, A., Röper, J.-C., Jülicher, F., and Eaton, S. (2010). Cell flow reorients the axis of planar polarity in the wing epithelium of Drosophila. Cell, 142(5):773–786.

Chan, L. N. and Gehring, W. (1971). Determination of blastoderm cells in Drosophila melanogaster. Proceedings of the National Academy of Sciences of the United States of America, 68(9):2217–2221.

Chen, Z., Medina, F., Liu, M.-y., Thomas, C., Sprang, S. R., and Sternweis, P. C. (2010). Activated RhoA binds to the pleckstrin homology (PH) domain of PDZ-RhoGEF, a potential site for autoregulation. The Journal of Biological Chemistry, 285(27):21070–21081.

Christie, J. M., Corchnoy, S. B., Swartz, T. E., Hokenson, M., Han, I.-S., Briggs, W. R., and Bogomolni, R. A. (2007). Steric interactions stabilize the signaling state of the LOV2 domain of phototropin 1. Biochemistry, 46(32):9310–9319.

Costa, M., Wilson, E. T., and Wieschaus, E. (1994). A putative cell signal encoded by the folded gastrulation gene coordinates cell shape changes during Drosophila gastrulation. Cell, 76(6):1075–1089.

Dawes-Hoang, R. E., Parmar, K. M., Christiansen, A. E., Phelps, C. B., Brand, A. H., and Wieschaus, E. F. (2005). folded gastrulation, cell shape change and the control of myosin localization. Development, 132(18):4165–4178.

Doubrovinski, K., Tchoufag, J., and Mandadapu, K. (2018). A simplified mechanism for anisotropic constriction in Drosophila mesoderm. Development, 145(24).

Fox, D. T. and Peifer, M. (2007). Abelson kinase (Abl) and RhoGEF2 regulate actin organization during cell constriction in Drosophila. Development, 134(3):567–578.

Gao, G.-J. J., Holcomb, M. C., Thomas, J. H., and Blawzdziewicz, J. (2016). Embryo as an active granular fluid: stress-coordinated cellular constriction chains. Journal of physics. Condensed matter : an Institute of Physics journal, 28(41):414021.

Gibson, D. G., Young, L., Chuang, R.-Y., Venter, J. C., Hutchison, C. A., and Smith, H. O. (2009). Enzymatic assembly of DNA molecules up to several hundred kilobases. Nature Methods, 6(5):343–345.

Gilmour, D., Rembold, M., and Leptin, M. (2017). From morphogen to morphogenesis and back. Nature, 541(7637):311–320.

Guntas, G., Hallett, R. A., Zimmerman, S. P., Williams, T., Yumerefendi, H., Bear, J. E., and Kuhlman, B. (2015). Engineering an improved light-induced dimer (iLID) for controlling the localization and activity of signaling proteins. Proceedings of the National Academy of Sciences, 112(1):112–117.

Han, K. (1996). An efficient DDAB-mediated transfection of Drosophila S2 cells. Nucleic Acids Research, 24(21):4362–4363.

Ip, Y. T., Park, R. E., Kosman, D., Yazdanbakhsh, K., and Levine, M. (1992). dorsal-twist interactions establish snail expression in the presumptive mesoderm of the Drosophila embryo. Genes & Development, 6(8):1518–1530.

Izquierdo, E., Quinkler, T., and De Renzis, S. (2018). Guided morphogenesis through optogenetic activation of Rho signalling during early Drosophila embryogenesis. Nature Communications, 9(1):2366.

Jaiswal, M., Dvorsky, R., and Ahmadian, M. R. (2013). Deciphering the molecular and functional basis of Dbl family proteins: a novel systematic approach toward classification of selective activation of the Rho family proteins. Journal of Biological Chemistry, 288(6):4486–4500.

Jiang, J., Kosman, D., Ip, Y. T., and Levine, M. (1991). The dorsal morphogen gradient regulates the mesoderm determinant twist in early Drosophila embryos. Genes & Development, 5(10):1881–1891.

Kerridge, S., Munjal, A., Philippe, J.-M., Jha, A., de las Bayonas, A. G., Saurin, A. J., and Lecuit, T. (2016). Modular activation of Rho1 by GPCR signalling imparts polarized myosin II activation during morphogenesis. Nature Cell Biology, 18(3):261–270.

Klueg, K. M., Alvarado, D., Muskavitch, M. A. T., and Duffy, J. B. (2002). Creation of a GAL4/UAS-coupled inducible gene expression system for use in Drosophila cultured cell lines. genesis, 34(1-2):119–122.

Ko, C. S. and Martin, A. C. (2020). The cellular and molecular mechanisms that establish the mechanics of Drosophila gastrulation. Current topics in developmental biology, 136:141–165.

Kölsch, V., Seher, T., Fernandez-Ballester, G. J., Serrano, L., and Leptin, M. (2007). Control of Drosophila gastrulation by apical localization of adherens junctions and RhoGEF2. Science, 315(5810):384–386.

Leptin, M., Casal, J., Grunewald, B., and Reuter, R. (1992). Mechanisms of early Drosophila mesoderm formation. Development (Cambridge, England). Supplement, pages 23–31.

Leptin, M. and Grunewald, B. (1990). Cell shape changes during gastrulation in Drosophila. Development, 110(1):73–84.

Manning, A. J., Peters, K. A., Peifer, M., and Rogers, S. L. (2013). Regulation of epithelial morphogenesis by the G protein-coupled receptor mist and its ligand fog. Science Signaling, 6(301):ra98–ra98.

Martin, A. C., Gelbart, M., Fernandez-Gonzalez, R., Kaschube, M., and Wieschaus, E. F. (2010). Integration of contractile forces during tissue invagination. The Journal of Cell Biology, 188(5):735–749.

Martin, A. C., Kaschube, M., and Wieschaus, E. F. (2009). Pulsed contractions of an actin-myosin network drive apical constriction. Nature, 457(7228):495–499.

Mason, F. M., Xie, S., Vasquez, C. G., Tworoger, M., and Martin, A. C. (2016). RhoA GTPase inhibition organizes contraction during epithelial morphogenesis. The Journal of Cell Biology, 127:jcb.201603077.

Medina, F., Carter, A. M., Dada, O., Gutowski, S., Hadas, J., Chen, Z., and Sternweis, P. C. (2013). Activated RhoA is a positive feedback regulator of the Lbc family of Rho guanine nucleotide exchange factor proteins. The Journal of Biological Chemistry, 288(16):11325–11333.

Morisato, D. and Anderson, K. V. (1995). Signaling pathways that establish the dorsal-ventral pattern of the Drosophila embryo. Annual Review of Genetics, 29(1):371–399.

Munjal, A., Philippe, J.-M., Munro, E., and Lecuit, T. (2015). A self-organized biomechanical network drives shape changes during tissue morphogenesis. Nature, 524(7565):351–355.

Nikolaidou, K. K. and Barrett, K. (2004). A Rho GTPase signaling pathway is used reiteratively in epithelial folding and potentially selects the outcome of Rho activation. Current Biology, 14(20):1822–1826.

Oakes, P. W., Wagner, E., Brand, C. A., Probst, D., Linke, M., Schwarz, U. S., Glotzer, M., and Gardel, M. L. (2017). Optogenetic control of RhoA reveals zyxin-mediated elasticity of stress fibres. Nature Communications, 8:15817.

Padash Barmchi, M., Rogers, S., and Häcker, U. (2005). DRhoGEF2 regulates actin organization and contractility in the Drosophila blastoderm embryo. The Journal of Cell Biology, 168(4):575–585.

Parks, S. and Wieschaus, E. (1991). The Drosophila gastrulation gene concertina encodes a G alpha-like protein. Cell, 64(2):447–458.

Piekny, A. J. and Glotzer, M. (2008). Anillin Is a Scaffold Protein That Links RhoA, Actin, and Myosin during Cytokinesis. Current Biology, 18(1):30–36.

Rauzi, M., Krzic, U., Saunders, T. E., Krajnc, M., Ziherl, P., Hufnagel, L., and Leptin, M. (2015). Embryo-scale tissue mechanics during Drosophila gastrulation movements. Nature Communications, 6(1):8677.

Sawyer, J. K., Harris, N. J., Slep, K. C., Gaul, U., and Peifer, M. (2009). The Drosophila afadin homologue Canoe regulates linkage of the actin cytoskeleton to adherens junctions during apical constriction. The Journal of Cell Biology, 186(1):57–73.

Schindelin, J., Arganda-Carreras, I., Frise, E., Kaynig, V., Longair, M., Pietzsch, T., Preibisch, S., Rueden, C., Saalfeld, S., Schmid, B., Tinevez, J.-Y., White, D. J., Hartenstein, V., Eliceiri, K., Tomancak, P., and Cardona, A. (2012). Fiji: an open-source platform for biological-image analysis. Nature Methods, 9(7):676–682.

Spahn, P., Ott, A., and Reuter, R. (2012). The PDZ-GEF protein Dizzy regulates the establishment of adherens junctions required for ventral furrow formation in Drosophila. Journal of Cell Science, 125(Pt 16):3801–3812.

Strickland, D., Lin, Y., Wagner, E., Hope, C. M., Zayner, J., Antoniou, C., Sosnick, T. R., Weiss, E. L., and Glotzer, M. (2012). TULIPs: tunable, light-controlled interacting protein tags for cell biology. Nature Methods, 9(4):379–384.

Sweeton, D., Parks, S., Costa, M., and Wieschaus, E. (1991). Gastrulation in Drosophila: the formation of the ventral furrow and posterior midgut invaginations. Development, 112(3):775–789.

Wagner, E. and Glotzer, M. (2016). Local RhoA activation induces cytokinetic furrows independent of spindle position and cell cycle stage. The Journal of Cell Biology, 213(6):641–649.

Weng, M. and Wieschaus, E. (2016). Myosin-dependent remodeling of adherens junctions protects junctions from Snail-dependent disassembly. The Journal of Cell Biology, 212(2):219–229.

Wenzl, C., Yan, S., Laupsien, P., and Großhans, J. (2010). Localization of RhoGEF2 during Drosophila cellularization is developmentally controlled by Slam. Mechanisms of Development, 127(7-8):371–384.

Yevick, H. G., Miller, P. W., Dunkel, J., and Martin, A. C. (2019). Structural Redundancy in Supracellular Actomyosin Networks Enables Robust Tissue Folding. Developmental Cell, 50(5):586–598.e3.

Zhang, Y., Werling, U., and Edelmann, W. (2012). SLiCE: a novel bacterial cell extract-based DNA cloning method. Nucleic Acids Research, 40(8):e55.

Zitserman, D. and Roegiers, F. (2011). Live-cell imaging of sensory organ precursor cells in intact Drosophila pupae. Journal of Visualized Experiments, (51).

